# Mitochondrial redox homeostasis links organellar stress surveillance to germline and somatic integrity in *Caenorhabditis elegans*

**DOI:** 10.64898/2025.12.03.691999

**Authors:** Marina Valenzuela-Villatoro, Patricia de la Cruz-Ruiz, David Guerrero-Gómez, Eva Gómez-Orte, Alfonso Schiavi, Silvia Maglioni, Mayte Montero, Rosalba I Fonteriz, José Carlos Casas-Martínez, Nicole E. Briand, Jingxiu Xu, María Jesús Rodriguez-Palero, Marta Artal-Sanz, Katarzyna Olek, Justyna Polaczyk, Michal Turek, Wojciech Pokrzywa, Suhong Xu, Javier E. Irazoqui, Brian McDonagh, Javier Álvarez, María Olmedo, Natascia Ventura, Juan Cabello, Antonio Miranda-Vizuete

**Affiliations:** Redox Homeostasis Group, Instituto de Biomedicina de Sevilla, IBiS/Hospital Universitario Virgen del Rocío/CSIC/Universidad de Sevilla, Seville, Spain; Centro de Investigación Biomédica de la Rioja (CIBIR), Logroño, La Rioja, Spain; IUF-Leibniz Research Institute for Environmental Medicine, Duesseldorf, Germany; Institute for Clinical Chemistry and Laboratory Diagnostic, Medical Faculty, Heinrich Heine University, Duesseldorf, Germany; Departamento de Bioquímica y Biología Molecular y Fisiología, Facultad de Medicina, Universidad de Valladolid, Valladolid, Spain; Unidad de Excelencia Instituto de Biomedicina y Genética Molecular (IBGM), Universidad de Valladolid y Consejo Superior de Investigaciones Científicas (CSIC), Valladolid, Spain; Discipline of Physiology, School of Pharmacy and Medical Sciences, University of Galway, Galway, Ireland; Apoptosis Research Centre, University of Galway, Galway, Ireland; Galway RNA Research Cluster, University of Galway, Galway, Ireland; Department of Microbiology, UMass Chan Medical School, Worcester, MA, USA; International Biomedicine-X Research Center of the Second Affiliated Hospital, Zhejiang University School of Medicine and the Zhejiang University-University of Edinburgh Institute, Haining, Zhejiang, China; Center for Stem Cell and Regenerative Medicine and Department of Burn and Wound Repair of the Second Affiliated Hospital, Zhejiang University School of Medicine, Hangzhou, Zhejiang, China; School of Basic Medical Sciences, Zhejiang University School of Medicine, Hangzhou, Zhejiang, China; Andalusian Centre for Developmental Biology, Consejo Superior de Investigaciones Científicas (CSIC), Universidad Pablo de Olavide, Junta de Andalucía. Department of Molecular Biology and Biochemical Engineering, Universidad Pablo de Olavide, Seville, Spain; Departamento de Ciencias de la Salud y Biomedicina, Facultad de Ciencias de la Salud, Universidad Loyola Andalucía. Dos Hermanas, Sevilla, Spain; Laboratory of Protein Metabolism, International Institute of Molecular and Cell Biology in Warsaw, Warsaw, Poland; Laboratory of Animal Molecular Physiology, Institute of Biochemistry and Biophysics, Polish Academy of Sciences, Warsaw, Poland; Departamento de Genética, Facultad de Biología, Universidad de Sevilla, Avenida Reina Mercedes s/n, 41012, Seville, Spain; Institute of Cell Biology, Heinrich Heine University Düsseldorf, Universitaetsstrasse 1, Duesseldorf; Department for the Promotion of Human Science and Quality of Life, University of Rome San Raffaele, Via di Val Cannuta 247, Rome, Italy

**Keywords:** mitochondria, elegans, thioredoxin reductase, glutathione reductase, unfolded protein response, ATFS-1

## Abstract

Mitochondrial redox homeostasis is essential for cellular metabolism and organismal development. To investigate the consequences of disrupting redox homeostasis in this organelle in a metazoan organism, we generated a double mutant lacking mitochondrial glutathione reductase (*gsr-1a*) and thioredoxin reductase (*trxr-2*) genes in *Caenorhabditis elegans*. While *gsr-1a* or *trxr-2* single mutants are phenotypically normal, double *gsr-1a trxr-2* mutants displayed small body size, gonadal migration defects, reduced brood size, and prolonged egg-laying period, without developmental delay or lethality. Transcriptomic analysis revealed strong induction of ATFS-1-dependent stress and detoxification genes. Consistent with this, *gsr-1a trxr-2* worms exhibited constitutive ATFS-1 nuclear localization and robust *Phsp-6::gfp* expression. Triple *gsr-1a trxr-2; atfs-1* mutants were nonviable, demonstrating that unfolded protein response (UPR^mt^) activation is essential under mitochondrial redox stress.

Despite the induction of a stress response at the transcriptional level, *gsr-1a trxr-2* double mutants were not more resistant to oxidative or pathogen stressors. Moreover, these mutants maintained normal respiration, ATP and ROS production while displaying altered mitochondrial morphology in a tissue-specific manner, independent of mitophagy genes but dependent on mitochondrial fission or fusion machinery. Functionally, *gsr-1a trxr-2* mutants showed impaired motility, reduced calcium uptake upon carbachol stimulation, enhanced hypodermal wound repair, and decreased fertilization efficiency associated with lower muscle exopher production.

Overall, our data show that simultaneous loss of mitochondrial GSR-1a and TRXR-2 compromises growth, fertility and muscle performance and triggers a constitutive ATFS-1-dependent UPR^mt^ that sustains viability revealing mitochondrial redox control as a core determinant of organismal proteostasis.

**Highlights:**

- *gsr-1a* or *trxr-2* single mutants have no overt phenotypes.
- *gsr-1a trxr-2* double mutants are viable but show small size, gonad migration defects and reduced progeny.
- Loss of both reductases in mitochondria triggers a constitutive ATFS-1–dependent UPR^mt^.
- ATFS-1 is essential for *gsr-1a trxr-2* worms survival.
- *gsr-1a trxr-2* animals remodel mitochondrial morphology in a tissue-specific manner.
- *gsr-1a trxr-2* double mutants exhibit impaired muscle and sperm function but enhanced wound healing.

**Graphical abstract (to be incorporated):** 

## Introduction

Mitochondria are subcellular organelles that not only play key roles in cellular function such as ATP synthesis, maintenance of Ca^2+^ homeostasis, Fe-S cluster biogenesis, heat production or lipid synthesis but are also a central hub for intracellular communication either by releasing signaling molecules or by direct interaction with other organelles to cope with cellular stress and metabolic demands (Monzel et al. 2023). To coordinate all these functions, mitochondria have developed a remarkable plasticity to form dynamic organellar tubular networks that are exquisitely controlled by fusion and fission mechanisms (Dorn 2019).

When mitochondrial function is compromised, for example by respiration inhibition, mitochondrial DNA damage, impairment of protein import or proteotoxic stress within the mitochondrial matrix, a protective response is initiated to counteract the insult, that mainly operates through two primary defense mechanisms. The first one acts at the molecular level and consists on a retrograde signal from mitochondria to the nucleus leading to the coordinated expression of a network of mitochondrial proteases, chaperones and membrane transporters, collectively known as the mitochondrial unfolded protein response (UPR^mt^), which is orchestrated by the transcription factor ATF5 in mammals or ATFS-1 in the nematode *C. elegans* (Fiorese et al. 2016; Nargund et al. 2012; Shpilka and Haynes 2018). When UPR^mt^ fails to cope with the mitochondrial insult, a second defense mechanism at the complete organelle level is triggered, by which the mitochondrial network is reorganized by fusion and fission mechanisms and mitophagy is activated to remove irreversibly damaged mitochondria with the ultimate objective of maintaining a healthy mitochondrial population (Picca et al. 2023; Tabara et al. 2025). Notably, in some cell types such as neurons and muscle cells, damaged mitochondria may also be extruded from the cell in large extracellular vesicles known as exophers, providing an additional route for clearing dysfunctional organelles beyond canonical mitophagy (Nicolas-Avila et al. 2020; Melentijevic et al. 2017; Turek et al. 2021).

Because mitochondria are the main site of reactive oxygen species (ROS) production, the mitochondrial redox status needs to be tightly controlled to prevent macromolecular damage such as oxidative misfolding by excessive ROS production (oxidative distress). At the same time, mitochondria must maintain a balanced redox-dependent signaling network to ensure proper intracellular communication and specific posttranslational modifications (oxidative eustress) (Sies et al. 2024; Sies et al. 2017). For this purpose, mitochondria are equipped with dedicated ROS detoxifying enzymes such as superoxide dismutases, glutathione peroxidases or peroxiredoxins but at the same time they also have enzymatic networks to regulate the redox balance like the thioredoxin and glutathione/glutaredoxin systems which use NADPH as source of reducing power (Jacobs and Riemer 2023; Hanschmann et al. 2013; Sies et al. 2017).

While initially considered as separate pathways, the cooperation between the thioredoxin and glutathione/glutaredoxin systems is now well established across species. For instance, in yeast lacking glutathione reductase, oxidized glutathione (GSSG) reduction is carried out by the cytoplasmic thioredoxin system composed of one thioredoxin reductase and two thioredoxins (Muller 1996; Tan et al. 2010). In *C. elegans*, cytoplasmic thioredoxin reductase TRXR-1 is functionally redundant with glutathione reductase GSR-1 to allow a proper molting progression (Stenvall et al. 2011). However, TRXR-1 is unable to rescue the embryonic lethality of *gsr-1* null mutants (Mora-Lorca et al. 2016), demonstrating developmental or tissue-specific requirements for this redundancy in the cytoplasm. In mice, mutations in the genes encoding the components of the cytoplasmic and mitochondrial thioredoxin systems result in embryonic lethal phenotypes (Conrad et al. 2004; Jakupoglu et al. 2005; Nonn et al. 2003; Matsui et al. 1996) while mutant mice lacking glutathione reductase are viable with very mild phenotypes (Rogers et al. 2004). Evidence for a functional redundancy between the two systems in the cytoplasm arises from studies in mice hepatocytes lacking both thioredoxin reductase and glutathione reductase that remain viable by boosting glutathione (GSH) *de novo* synthesis through the transsulfuration pathway (Eriksson et al. 2015; Prigge et al. 2012). This functional redundancy is not restricted to animals but also includes plants, as *Arabidopsis thaliana* defective in both cytoplasmic thioredoxin reductase NTRA and glutathione reductase GR1 enzymes are not viable with a pollen lethal phenotype because of their inability to fertilize wild-type ovules (Marty et al. 2009).

In contrast with the functional redundancy of the cytoplasmic thioredoxin and glutathione systems, less is known on their cooperation within mitochondria. Indeed, to our knowledge, this redundancy has only been shown in yeast and *Arabidopsis thaliana*. Interestingly, in *Saccharomyces cerevisiae* and *Schizosaccharomyces pombe*, the absence of mitochondrial glutathione reductase causes a major disruption of cellular redox homeostasis which is only partly alleviated by the mitochondrial thioredoxin system (de Cubas et al. 2025; Gostimskaya and Grant 2016) while in *Arabidopsis thaliana* the mitochondrial thioredoxin system and the GSSG exporter ATM3 coordinate to back up the absence of mitochondrial glutathione reductase GR2 (Marty et al. 2019). However, it is not known whether this functional redundancy operates in metazoan.

To fill this gap, we have approached the question using the nematode *C. elegans*, a well-established model organism, taking advantage of its genetic tractability, high conservation of redox signaling pathways and its transparency throughout all its developmental stages, allowing *in vivo* imaging approaches (Corsi et al. 2015). In this work, we investigate whether the mitochondrial thioredoxin and glutathione systems exhibit functional redundancy in a metazoan and show that concurrent loss of mitochondrial *trxr-2* and *gsr-1a* disrupts redox homeostasis and elicits a strong ATFS-1–dependent UPR^mt^ that is essential for organismal viability. Double mutants exhibit reduced body size, defects in gonadal migration, and compromised fertility, revealing key physiological processes sensitive to mitochondrial redox imbalance. Moreover, we uncover tissue-specific alterations in mitochondrial dynamics that impair muscle performance, hypodermal repair responses, and sperm function. Together, these findings establish mitochondrial redox control as a critical determinant of cellular and organismal proteostasis.

## Materials and methods

### C. elegans strains

The standard methods used for culturing and maintenance of *C. elegans* were as previously described (Stiernagle 2006). A list of all strains used and generated in this study is provided in Supplementary Table 1. The *gsr-1a(syb266)* allele was generated at SunyBiotech (https://www.sunybiotech.com/) by CRISPR-Cas9 editing. All VZ strains are 6x outcrossed with N2 wild type, except those strains generated by CRISPR-Cas9 (*syb* alleles) that were 2x outcrossed. Worm reagents and details on the protocols used for genotyping the different alleles reported in this work can be provided upon request. Unless otherwise noted, all experiments were performed on synchronized worms generated by allowing 10 to 15 gravid hermaphrodites to lay eggs during two to three hours on seeded plates at 20°C.

### Size, gonad migration defects and oocyte quantification

Quantification was carried out in animals grown for 72 h at 20 °C after synchronized egg-lay. Worms were paralyzed with 10 mM levamisole (Sigma-Aldrich, Cat. #L9756) on a microscope slide in an Olympus BX61 fluorescence microscope equipped with a DP72 digital camera coupled to CellSens Software. Images were acquired and the length and area of the animals were determined using ImageJ software (Schneider et al. 2012). Quantification of gonad migration defects and the number of oocytes was directly performed on immobilized animals at the microscope. The number of oocytes corresponds to the proximal arm of the posterior gonad.

### Developmental timing

We measured developmental timing of single worms using a bioluminescence-based method previously described (Olmedo et al. 2015) but using the reporter *sevIs1 [Psur-5::luc+::GFP]* (Rodriguez-Palero et al. 2018). To obtain age-synchronized eggs, we transferred 10-15 hermaphrodites to a fresh NGM plate and allowed them to lay eggs for one hour. Synchronized eggs were individually placed in wells of a 96 well-plate containing 100 μl of S-basal plus 10 μg/ml cholesterol (Sigma-Aldrich, Cat. #C8667) with 200 μM D-Luciferin (Biothema BT11100). After placing all embryos, 100 μl of S-basal with 20 g/l *E. coli* OP50-1 was added to each well. Samples from different strains were alternated on the plate to avoid local effects. Plates were sealed with a gas-permeable membrane and transferred to a luminometry reader (Berthold Centro XS3) placed inside a cooled incubator (Panasonic MIR-254). Luminescence was recorded for 1 s per well, at 5-min intervals.

### Embryonic arrest and brood size quantification

For embryonic arrest, 10 gravid hermaphrodites were allowed to lay eggs for one hour in three replica plates. After removal of the parents, the embryos were counted and allowed to develop three days at 20 °C. Viable progeny was quantified and detracted to the number of laid embryos to determine the embryonic arrest. Brood size was quantified from 20 L4 hermaphrodites, placed in groups of five animals on four separate plates. Worms were transferred to fresh plates every 24 hours, and the number of viable progeny was recorded daily for a total of eight days. The total brood size was calculated as the sum of all viable progeny by each worm throughout the experimental period.

### RNAseq and q-PCR

Worms were synchronized using bleaching solution (50 % dH_2_O (v/v)), 30 % commercial NaClO (v/v)) and 20 % NaOH (5N) (v/v)) and grow until young adults. For total RNA extraction, young adult worms were lysed using mirVana Kit (Ambion, Cat. #1560), following manufactureŕs instructions. Homogenization of the lysate was performed using a conventional rotor-stator homogenizer polytron, pre-chilled with liquid nitrogen. Global gene expression profile was analyzed at the Genomic Platform of the CIBIR (http://cibir.es/es/plataformas-tecnologicas-y-servicios/genomica-y-bioinformatica). Expression analysis was performed as described (Gomez-Orte et al. 2018) using DESeq2 (Love et al. 2014) and edgeR (Robinson et al. 2010) algorithms.

For q-PCR analysis, RNA was treated with DNAse to eliminate any DNA contamination. In each sample, a total reaction of 10 µl contained: 500ng RNA, 1 µl RNase-Free DNAse (Promega, Cat. #M6101), 1 µl 10x Reaction Buffer and DEPC water. Reactions were incubated for 30 min at 37 °C and then stopped by adding 1 µl STOP solution (Promega, Cat. #M6101) and further incubation for 15 min at 65 °C. cDNA synthesis was performed using SuperScript III First-Strand Synthesis System for RT-PCR (Invitrogen, Cat. #18080051) following instructions for random hexamers primed. cDNA was eluted in TE buffer. q-PCR analysis Power SYBR Green Master Mix (ThermoFisher Scientific, Cat. #4367659) and specific primers were used in a QuantStudio 5 Real Time PCR System (Applied Biosystems, ThermoFisher). Normalization to actin *act-1* expression was used to calculate relative expression. The experiments were carried out in three independent replicas. Primers pairs (5’->3’) used were: *msp-76* Fw1: TGACAAGCACACCTACCACA, *msp-76* Rv1: TCCTTTGGGTCGAGAACTCC; *clec-4* Fw1: TCCACAATACCGCCTCAACT, *clec-4* Rv-1: CACGTCCGCCAAATGTCATA; *cyp-14A4* Fw1: TCACAAGCCACCAGAGTCAT, *cyp-14A4* Rv1: CGCGGAGGATTTTCAGTGAG; *ugt-19* Fw1: TCCAGTCGCCTTGATCATGT, *ugt-19* Rv1: TTGCCACATAGTCATCCGGT; *F22B3.7* Fw1: AAATGATGCTGGTATGGGGC, *F22B3.7* Rv1: CCTTCCAAGCCATCACACAC; *act-1* Fw1:CCAGGAATTGCTGATCGTATG, *act-1* Rv1: GGAGAGGGAAGCGAGGATAG

### Fluorescence and Light Microscopy Imaging

Brightfield, differential interference contrast (DIC), and fluorescence images were acquired using an Olympus BX61 fluorescence microscope equipped with a DP72 digital camera and CellSens software for image acquisition and analysis. First day adult worms were mounted on 2% agar pads and anesthetized with 10 mM levamisole prior imaging. For mitophagy quantification using the *Is [Pmyo-3::tomm-20::Rosella]* transgene (Palikaras et al. 2015), FITC and TRITC fluorescence filters were used sequentially to calculate the GFP/DsRed fluorescence intensity ratio at 20x magnification. Images were analysed using ImageJ Software (Schneider et al. 2012).

### Confocal Imaging

High-resolution confocal imaging was performed using a Nikon A1R confocal microscope equipped with a Plan Apo VC 60x/1.4 NA oil immersion objective. Image acquisition was conducted using the NIS-Elements software and the ND Acquisition module. All Animals were mounted on 2% agar pads and anesthetized with 10 mM levamisole prior to imaging except for mitochondrial structure.

For the analysis of mitochondrial structure in muscle cells using the *wacIs14[Pmyo-3::tomm-20_1-50aa::attB5::mgfp]* transgene (Turek et al. 2021)) and hypodermal tissue with the *juEx4796 [Pcol-19::mito::gfp; Pttx-3::rfp]* transgene (Xu and Chisholm 2014a)), first day adult worms were immobilized on 5% agarose pads using polystyrene nanoparticles (Alpha Aesar, polystyrene latex microspheres, 0.10 µm, 2.5% w/v dispersion in water, Cat. #42712). Muscle tissue was imaged using Z-stack acquisition followed by maximal intensity projection. For hypodermal tissue, a single focal plane was selected for analysis. Imaging was performed using the same microscope setup as described above. Automated image processing was carried out using a custom ImageJ macro developed by Alejandro Campoy (Microscopy Facility, CABD). The pipeline included background subtraction, median filtering, and automatic thresholding to generate binary images. Mitochondrial features, such as elongation, circularity, and the percentage of area occupied, were quantified according to previously described methods (Zhao et al. 2017).

To quantify ATFS-1 subcellular localization using the *mskEx1 [Patfs-1::atfs-1::gfp]* transgene (Nargund et al. 2012), full-body images of L4 larvae were acquired using a single focal plane and tiled XY scans were used to generate representative images. Imaging parameters were optimized to enable whole-organism reconstruction, capturing both fluorescence and differential interference contrast (DIC) channels. Worms that displayed obvious nuclear labeling in at least 2 intestinal cells were considered as positive for ATFS-1 nuclear localization.

For the quantification of sperm cells in the spermatheca with the *oxIs318 [Pspe-11::mCherry::histone]* transgene (Frokjaer-Jensen et al. 2008), sperm nuclei were imaged using Z-stack acquisition followed by maximal intensity projection and saved as 8 bits images. The sperms nuclei were quantify using ImageJ wand tool for each spermatheca.

For hyp7 nuclei quantification using the *madEx16 [Pdpy-7::4xNLS::GFP]* transgene (de Lucas et al. 2015), a 20x oil-immersion objective was used to image entire animals. Z-stack images were acquired and processed using maximum intensity projection to quantify nuclei from the posterior part of the animals on a single lateral side.

### MitoTracker Deep Red Assay

Mitochondrial staining was performed using MitoTracker™ Deep Red FM (Invitrogen, Cat. #M22426). A 1 mM stock solution was prepared in DMSO (Panreac, Cat. #131954), and the working solution was obtained by a 1:10 dilution in water to a final concentration of 100 µM, following the protocol described by Artal-Sanz and Tavernarakis (Artal-Sanz and Tavernarakis 2009). A volume of 40 µL of the working solution was applied to 60 mm × 15 mm plates. Plates were incubated in the dark for 1 hour to allow uniform diffusion of the dye, after which L4-stage worms were transferred and incubated for at least 16 hours at 20 °C. Following incubation, worms were immobilized on 5% agarose pads using polystyrene nanoparticles (Alpha Aesar, Cat. #42712) as described above. Image acquisition was performed within 30 minutes using the Nikon A1R confocal microscope with the 60×/1.4 NA objective and the Deep Red laser channel. Image processing and mitochondrial quantification were conducted using the same workflow described for tissue-specific imaging.

### TMRE/Mitotracker Red CMXRos and Green Staining and Imaging

To assess mitochondrial membrane potential, Tetramethylrhodamine, Ethyl Ester, Perchlorate (TMRE) (Invitrogen, Cat. #T669) dye was diluted to a concentration of 10 mM in DMSO, added to molten NGM medium to a 2.5 µM final concentration and poured into plates, which were then allowed to dry in the dark for 30 minutes. Plates were subsequently seeded with *E. coli* OP50 and incubated for 48 hours. L4-stage worms were then transferred onto the plates and incubated for at least 16 hours at 20 °C.

For the quantification of mitochondrial mass and reactive oxygen species (ROS), a combination of MitoTracker™ Green FM and MitoTracker™ Red CM-H_2_XRos were used (Invitrogen, Cat. #M7514 and #M7513, respectively). Both dyes were diluted to 1 mM in DMSO and a working solution was prepared in water at 10 µM. Mitotracker dyes were added onto 35 mm NGM plates previously seeded with *E. coli* OP50 to a 100 nM final concentration. After allowing the plates to dry for 30 minutes, L4-stage worms were transferred and incubated for a minimum of 16 hours at 20°C.

Prior to imaging for all mitochondrial dyes, worms were cleaned by crawling on unseeded NGM plates for 30 minutes (TMRE) and 60 min (Mitotrackers). Fluorescence (FITC or TRITC), brightfield, and differential interference contrast (DIC) images were acquired using an Olympus BX-61 upright microscope equipped with UPlanFl 10x/0.30 objective (U Plan Fluor series) and DP72 digital camera coupled to CellSens Software for image acquisition and analysis. Animals were mounted on 2% agar pads and anesthetized with 10 mM levamisole prior to imaging. The whole animals were quantified by ImageJ.

### Oxygen Consumption Rate (OCR) Measurement

The oxygen consumption rate (OCR) was measured using a Seahorse XFp Analyzer (Agilent Technologies). Approximately 25 *C. elegans* at day 1 adult stage were transferred into each well of Seahorse XFp cell culture miniplates pre-filled with M9 buffer. OCR was recorded at 20 °C, eight times under basal conditions, ten times after carbonyl cyanide 4-(trifluoromethocy)phenylhydrazone (FCCP) injection (for maximal respiration) and four times after sodium azide (for no mitochondrial respiration). Working solutions were prepared in M9 buffer at the following final concentrations: FCCP (Sigma-Aldrich, Cat. #C2920) at 250 μM and sodium azide (Sigma-Aldrich, Cat. #S2002) at 400 mM. Basal OCR was defined as the average of the last five initial measurements. Post-treatment OCR was calculated as the average of the last four measurements after FCCP injection and last two after sodium azide injection.Values were normalized to the number of worms per well. A minimum of three independent experiments were performed.

### ATP levels Measurement

ATP levels were measured according to the published protocol (Palikaras and Tavernarakis 2016), with minor modifications, as described below. For each assay, approximately 100 day 1 adult worms were used for ATP and total protein extraction. The resulting supernatants were diluted threefold to obtain a final volume of 150 µL. ATP concentrations were determined using a bioluminescence-based ATP Determination Kit PRO, 10 mL (Biaffin GmbH & Co., Cat. #LBR-P010), following the manufacturer’s instructions. Luminescence was measured with a CLARIOstar plate reader (BMG LabTech) using Invitrogen™ microplates designed for fluorescence-based assays (Cat. #M33089). ATP standard curves were prepared at concentrations of 5, 10, 15, and 25 µM.

### Staphylococcus aureus and Pseudomonas aeruginosa infection assay

Bacterial infection assays were performed using *Staphylococcus aureus* strain SH1000 *telA::KAN* and *Pseudomonas aeruginosa* strain PA14 following standard methods (Powell and Ausubel 2008), with minor modifications as detailed next. *S. aureus* was cultured in Tryptic Soy Broth (TSB; BD Biosciences, Cat# 211825) supplemented with 50 µg/ml kanamycin (Sigma-Aldrich, Cat# K1377) overnight (37 °C, 200–220 rpm), and 10 µl was spread on Tryptic Soy Agar plates (TSA; BD Biosciences, Cat# 236950) containing 10 µg/ml kanamycin, incubated at 37 °C for 5 h, and shifted to 25 °C overnight. *P. aeruginosa* was cultured overnight in Luria-Bertani broth (LB; BD Biosciences, Cat# 244620), and 10 µl was spread on Nematode Growth Media (NGM) slow killing plates [prepared using Bacto Agar (BD Biosciences, Cat# 214010) and Bacto Peptone (BD Biosciences, Cat# 211677)], incubated at 37 °C for 24 h, and shifted to 25 °C for 24 h to obtain full bacterial lawns. *C. elegans* were grown to the L4 larval stage on *E. coli* OP50 at 20°C. For both assays, L4 animals were treated with 100 µg/ml 5-fluoro-2′-deoxyuridine (FUDR; Sigma-Aldrich, Cat# F0503) on solid media overnight at 15 °C. Animals were subsequently transferred to infection plates lacking FUDR. Three plates each containing 30–40 animals were examined for each genotype. Infection assays were carried out at 25 °C. Survival was quantified using standard methods (Powell and Ausubel 2008); animals dying of disrupted vulva or crawling on walls were censored for analysis.

### Paraquat and Juglone tolerance assay

Juglone (Sigma-Aldrich, Cat. #H47003) and paraquat (Sigma-Aldrich, Cat. # 856177) were dissolved in DMSO and Milli-Q water and then added to molten NGM to obtain final concentrations of 150 µM (juglone) and 50 mM (paraquat). The media were thoroughly mixed and poured onto 60 mm (juglone) or 35 mm (paraquat) diameter plates. Once solidified, plates were seeded with 50 µL of *E. coli* OP50 and allowed to dry before use. L4-stage worms were then transferred and incubated at 20 °C for 16 hours before scoring for live or dead animals. Worms were considered alive if they responded to gentle touch with a platinum pick on the agar surface or on the head or showed pharyngeal pumping. Worms desiccated on the wall of the plate were censored and excluded from the analysis.

### Rotenone and Antimycin A tolerance assay

Rotenone (Sigma-Aldrich, Cat. #R8875) and antimycin A (Sigma-Aldrich, Cat. #A8674) mitochondrial electron transport chain (ETC) inhibitors, were dissolved in DMSO to a stock solution of 5 mg/ml and 10 mg/ml and then added to molten NGM medium to final concentrations of 1 µM and 20 µM, respectively. Once solidified, plates were seeded with 50 µL of *E. coli* OP50 and allowed to dry before use. L4-stage worms were then transferred and incubated at 20°C for 16 hours before scoring for live or dead animals. Worms were considered alive if they responded to gentle touch with a platinum pick on the agar surface or on the head or showed pharyngeal pumping. Worms desiccated on the wall of the plate were censored and excluded from the analysis.

### Lifespan assays

Lifespan analyses were carried out with standard procedure in the field (Schiavi et al. 2015). Briefly, age-synchronized population of 60-80 worms were used to start survival analysis. To avoid cross generation contamination worms were transferred to fresh plates every day during the fertile phase while every other day from the end of the fertile period. Animals not able to move upon pick-prodding and with no pharyngeal pumping were scored as dead. Survival analysis was performed with OASIS 2 (Han et al. 2016) using the Kaplan Meier estimator. Statistical differences were evaluated using the log-rank test between the pooled population of worms and p-values were adjusted for multiple comparisons by Bonferoni method.

### Embryonic and larval lethality assay

To quantify embryonic and larval lethality, 15–20 synchronized hermaphrodites were allowed to lay eggs for 2-3 hours and laid embryos were determined. After three days of incubation at 20 °C the developmental stages of the progeny were quantified in a stereomicroscope.

### CeleST Swimming activity assay

Swimming behavior was assessed at day 1 of adulthood. A glass slide with a 10mm pre-printed ring was prepared and loaded with 50μl M9 Buffer. Five worms were placed in the swimming area, for a total of 45 animals per condition. Worms were allowed to settle for 20s before starting the recording. 30s videos with at 16 frames per second of the animals were taken using a Nikon LV-TV microscope with a OPTIKA C–P20CM camera and processed by CeleST software (Restif et al. 2014).

### Pharyngeal pumping

Pharyngeal pumping measurements were performed by recording 20-second videos for each first day adult worm using a Leica MDG36 fluorescence stereo microscope equipped with a DMK 31AU03 monochrome camera. Video acquisition was carried out with IC Capture 2.3 software. Pharyngeal pumps were manually counted using video playback at 0.42× speed. Final pumping rates were converted to units of pumps per minute through appropriate time normalization.

### Electropharyngeogram

To perform electrical recordings of pharyngeal pumping (electropharyngeograms, or EPGs), we placed the Nemametrix Screen Chip System on an inverted Zeiss Axiovert 200 microscope equipped with an LD A-Plan 20x objective. To minimize interfering electrical noise during the recordings, we used a system grounding shield. Baseline noise was typically between 10 and 40 µV. The experiments were performed at 20°C. For each experiment, 100 worms at day 4 of adulthood were picked from the culture plate and washed in 1.5 ml of 0.2 µm filtered M9 buffer containing 0.1% Tween. The worms were then washed four times with 0.2 µm filtered M9 buffer, once with M9 buffer containing 2.3 mM serotonin (Aesar Chemicals, Cat. #B21263), and finally suspended in 1 ml of M9 buffer containing 2.3 mM serotonin. They were then left to settle for 15 minutes. All experiments were performed within 15–90 minutes of initial serotonin exposure. We loaded a NemaMetrix Screen Chip System (NemaMetrix, Eugene, OR, USA; Cat. #SK100) with a fresh SC40 Screen Chip (NemaMetrix, Eugene, OR, USA; Cat. #SKU0002) and added an M9 buffer solution containing 2.3 mM serotonin. After initiating the NemAcquire software and recording the basal power line noise, the worms were vacuum-loaded onto the SC40 Screen Chip to start the experiment. The 1 Hz high-pass and 50 Hz notch filter settings were selected. An EPG recording lasting 180 seconds was made for each animal, and records from 20–25 animals were obtained for each replicate. These records were analysed using NemAnalysis v0.2 software. Experiments with a frequency of less than 0.1 Hz or a pump duration coefficient of variation greater than 50% were censored.

### Mitochondrial calcium imaging

Measurements of mitochondrial Ca^²⁺^ in the pharynx were performed in worms expressing the *Ex [Pmyo-2::2mt8::YC3.60]* reporter (Alvarez-Illera et al. 2017) worms at day 5 of adult life. The worms were glued onto an agar pad (2% agar in M9 buffer) on a coverslip using Dermabond Topical Skin Adhesive (Johnson & Johnson), and 2.3 mM serotonin (Aesar Chemicals, Cat. #B21263) was added to stimulate pumping. The coverslip containing the glued worm was mounted in a chamber on the stage of a Zeiss Axiovert 200 inverted microscope. Fluorescence was excited at 430 nm using a Cairn monochromator with a 15 nm bandwidth, and images of the emitted fluorescence were obtained using a Zeiss C-Apochromat 40x/1.2 W objective. These images were collected using a 450 nm long-pass dichroic mirror and a Cairn Optosplit II emission image splitter, which produced separate emission images at 480 nm and 535 nm. The splitter contained DC/ET480/40m and DC/ET535/30m emission filters and a FF509-FDi01-25×36 dichroic mirror (all from Chroma Technology). A Hamamatsu ORCA-ER camera recorded simultaneous 200 ms images at the two emission wavelengths continuously at a rate of 2.5 Hz. Both images were then ratioed to obtain 535 nm/480 nm fluorescence ratio images of the pharynx terminal bulb. Experiments were performed at 20°C, with basal fluorescence being recorded for 15 minutes prior to the addition of carbachol to a final concentration of 10 mM to induce maximum mitochondrial Ca²_⁺_ uptake.

Fluorescence was recorded and analysed using the Metafluor programme (Universal Imaging). The traces shown represent either the 535 nm or 480 nm emission fluorescence, or the 535 nm/480 nm fluorescence emission ratio, of the pharynx terminal bulb. [Ca²_⁺_] peaks in the ratio were only considered acceptable when inverted changes at both wavelengths were clearly observed, as shown in Figure 6e-f. The fluorescence intensities and changes in the ratio were analysed using an algorithm designed to calculate the width at baseline and mid-height, both of which were expressed in seconds. The height was obtained as a percentage of the change in the ratio, and the frequency was measured at each peak by dividing 9 by the time interval including that peak and the four peaks before and after it. The mean value was calculated as an average of all the individual values obtained for each parameter.

### Hypodermal wounding

We wounded the epidermis of young adult stage animals using a Micropoint UV laser or needle. All images before and after wounding were taken using a spinning disk confocal microscope (Andor 100x, NA 1.46 objective). Actin ring quantitation and survival rate were performed as previously described (Xu and Chisholm 2014b). The nuclear intensity analyses were performed in ImageJ. Background intensity was used as a threshold. The intensity was analyzed using Analyze Particles (a function in ImageJ software) automatically with minimum size at 0.2 square micrometers. The value of intensity was automatically calculated and analyzed using GraphPad. At least 20 wounded young adult animals were examined. For cluster number, 600 × 400-pixel around the wounded area was chosen for analysis.

### Fertilization capacity

To assess male fertility, *fem-1(hc17)* hermaphrodites (grown at 25 °C) were crossed with wild-type or *gsr-1a trxr-2* young adult males in a 4:1 ratio. After 24 hours, males were removed and hermaphrodites were transferred to a new plate at 20 °C. The hermaphrodites were transferred daily until they stopped laying eggs. The total brood size was counted. Hermaphrodites that died or were absent before the end of the experiment were excluded from the analyses (Mei et al. 2023).

### Exophers

Exopher quantification followed the procedures outlined by Turek *et al*. (Turek et al. 2021) and (Banasiak et al. 2023). Briefly, exophers were assessed using a Zeiss Axio Zoom.V16 stereomicroscope equipped with 63 HE and 38 HE filter sets. Worms were age-matched by manually selecting 3-fold stage embryos under a stereomicroscope. On the second day of adulthood, animals were examined directly on NGM plates, and the number of exophers produced by each freely moving worm was counted.

### Graphical and Statistical Analysis

Data were processed in Excel (Microsoft Corporation) then Prism (GraphPad Software) was used to generate curve and bar charts and perform the statistical analyses described in the Figure Legends.

## Results

### Inactivation of mitochondrial glutathione reductase and thioredoxin reductase causes germline defects and small body size

GSH is the main redox currency in the cell and its key role in mitochondrial function is illustrated by the inability to proliferate of mammalian cells lacking GSH mitochondrial importers SLC25A39 and SLC25A40 (Wang et al. 2021). This essential role of GSH in mitochondria is conserved in *C. elegans* as a null allele of the only worm SLC25A39/SLC25A40 orthologue *C16C10.1(syb5556)* results in a fully penetrant L1 larval lethality and can only be propagated as a balanced heterozygous (strain VZ1095, Supplementary Table 1). Once inside mitochondria, GSH redox homeostasis is maintained by the redundant action of the mitochondrial thioredoxin and glutathione/glutaredoxin systems in organisms as distant as yeast or *Arabidopsis thaliana* (Gostimskaya and Grant 2016; Marty et al. 2019).

To test whether this functional redundancy is also present in metazoan, we investigated the phenotypic consequences of disrupting mitochondrial redox homeostasis in *C. elegans* by combining the previously characterized *trxr-2(ok2047)* null allele of mitochondrial thioredoxin reductase (Cacho-Valadez et al. 2012) (Fig 1a) with a newly generated CRISPR-Cas9 *gsr-1(syb266)* deletion allele, which removes the first exon of *gsr-1* (Fig 1a). This exon encodes the mitochondrial targeting sequence of GSR-1a isoform, so the *gsr-1(syb266)* allele permits expression of the essential cytosolic isoform GSR-1b, while eliminating expression of the mitochondrial isoform GSR-1a (Mora-Lorca et al. 2016).

**Figure 1.**
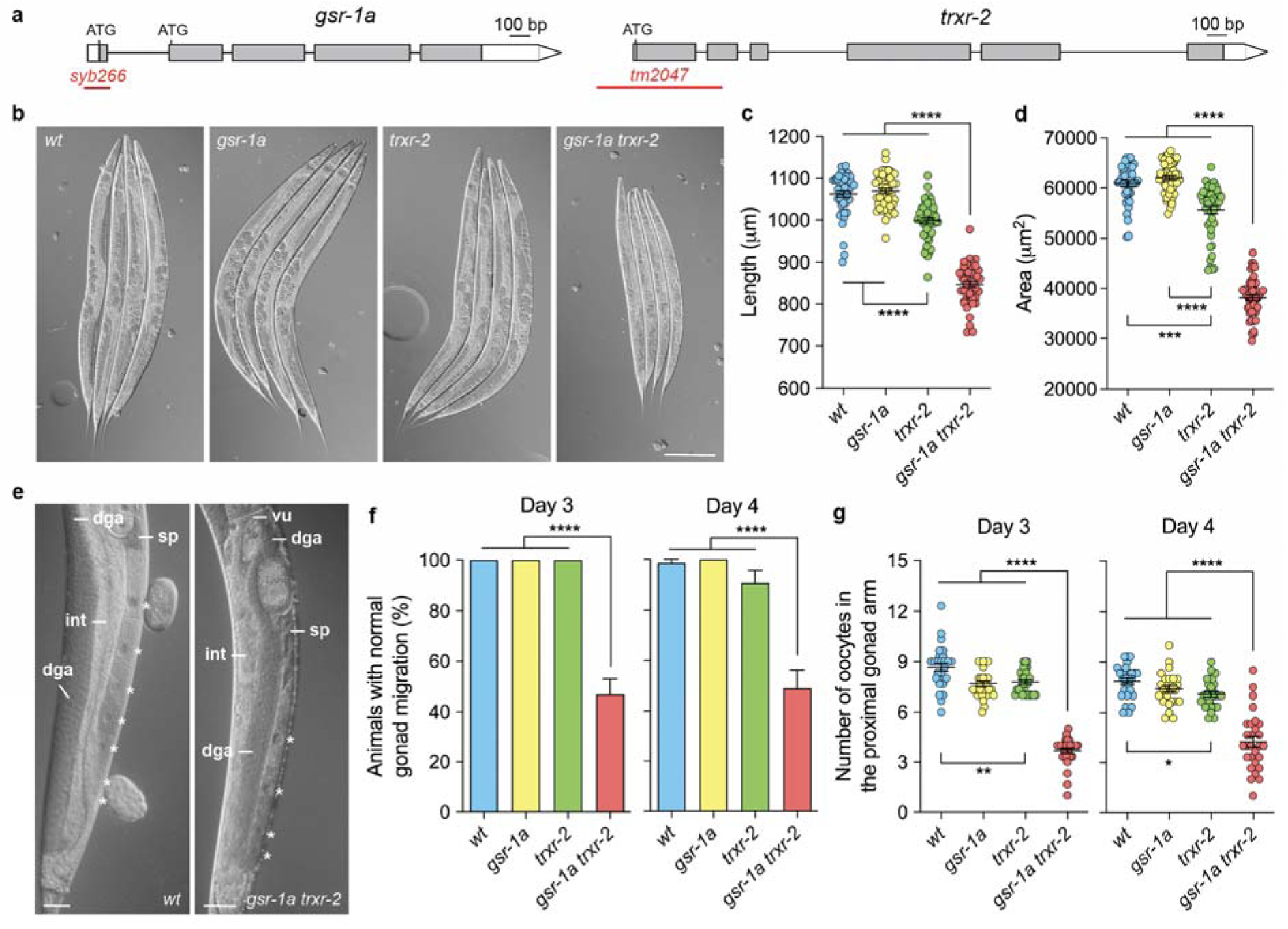
Genomic organization of *gsr-1* and *trxr-2* genes and visible phenotypes of *gsr-1a(syb266)* and *trxr-2(tm2047)* mutants. a) Boxes represent exons and lines show spliced introns. White boxes indicate 5’-UTR and 3’-UTR, respectively, and grey boxes indicate the ORF. Boundaries of *gsr-1a* and *trxr-2* deletions are shown as red lines. b) Representative micrographs of animals of the specified genotypes at third day after synchronized egg-lay. Scale bar 200 µm. Quantification of c) worm length and d) worm area. Data are from two independent experiments with at least 25 animals per assay. Error bars are SEM. ***p<0.001; ****p<0.0001 by Kruskal-Wallis test. e) Representative micrographs of wild-type controls and *gsr-1a trxr-2* double mutants at third day after synchronized egg-lay. dga, distal gonad arm; int, intestine; sp, spermatheca; vu, vulva. Scale bar 25 µm. f) Quantification of worms with gonad migration defects. Data are the mean +/− SEM from three different experiments with at least 20 animals per assay. ****p<0.0001 by Kruskal-Wallis test. g) Quantification of the oocytes in the proximal gonad arm. Data are from three different experiments with at least 20 animals per assay. Error bars are SEM. *p<0.05; ** p<0.01; **** p<0.0001 by Kruskal-Wallis test.

While the single mutants appeared superficially wild type with no overt phenotype, the *gsr-1a trxr-2* double mutants displayed a significantly smaller size, which was already evident at the L4 stage (two days at 20 °C after synchronized egg-lay) (Sup Fig 1), phenotype that became more pronounced by the first day of adulthood (Fig 1b-d). Additionally, we observed a high incidence of gonad migration defects, characterized by thinning and mislocalization of the distal gonad arm relative to the intestine, sometimes extending into the vulva area (Fig 1e,f). There was also reduced thickness of the proximal gonad arm, which contained fewer and slimmer oocytes, along with a distorted spermatheca (Fig 1e,g).

**Supplemental Figure 1.**
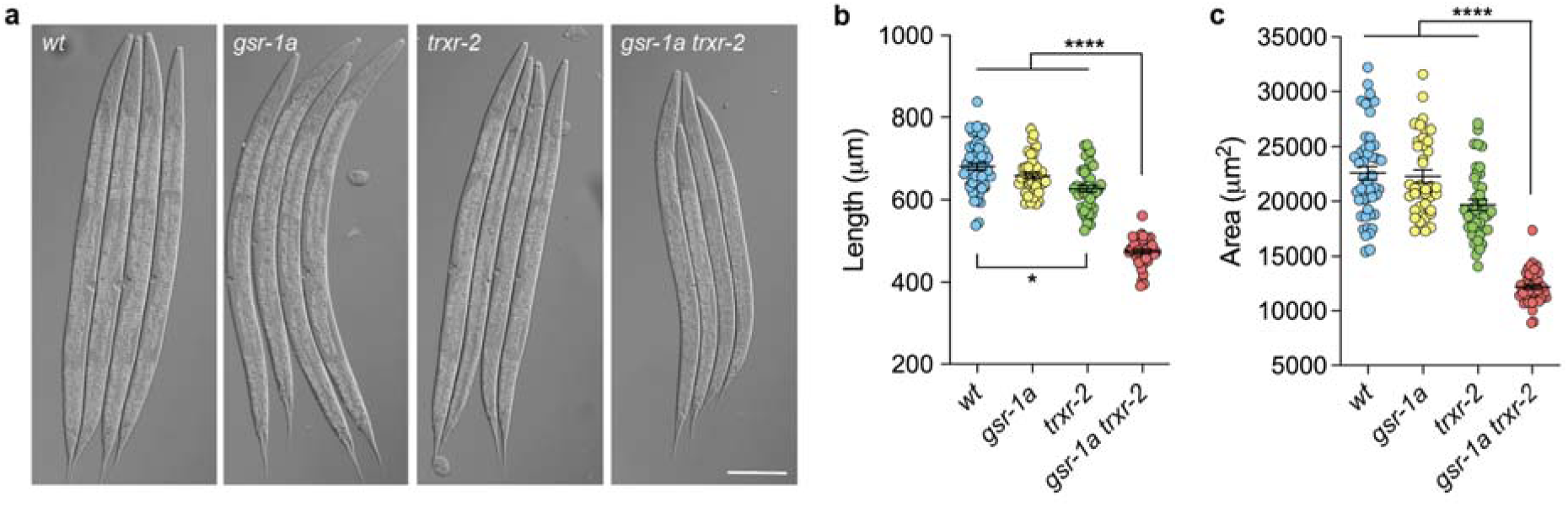
Size phenotypes of *gsr-1a(syb266)* and *trxr-2(tm2047)* mutants. a) Representative micrographs of animals of the specified genotypes at second day after synchronized egg-lay. Scale bar 100 µm. Quantification of b) worm length and c) worm area. Data are from two independent experiments with at least 25 animals per assay. Error bars are SEM. *p<0.05; **** p<0.0001 by Kruskal-Wallis test.

### *gsr-1a trxr-2* double mutants exhibit normal developmental timing but have reduced brood size and extended egg-lay period

Interestingly, during the isolation of the *gsr-1a trxr-2* double mutants, we observed that OP50 plates seeded with double mutant candidates became saturated noticeable later than those seeded with either single mutants or wild-type controls, suggesting a developmental delay and/or reduced progeny production in the double mutants, both phenotypes previously associated with mitochondrial dysfunction (Dillin et al. 2002; Curran et al. 2004; Artal-Sanz et al. 2003).

To directly assess whether the reduced plate saturation reflected a bona fide developmental delay, we quantified the duration of embryonic and larval stages in the double mutants using the luciferase-based method developed by Olmedo *et al*. (Olmedo et al. 2015) that accurately quantifies the duration of all molt and intermolt periods as well as the time from egg-lay to hatching and from the last molting step until the lay of the first embryos (Fig 2a). We found that the embryonic and larval development of *gsr-1a trxr-2* animals do not differ from that of wild type and *gsr-1a* or *trxr-2* single mutant controls (Fig 2b,c). However, we noticed that the period from the last molt until the first embryos are laid was significantly longer in *gsr-1a trxr-2* double mutants (Fig 2d) suggesting problems with progeny production caused by oocyte production, sperm depletion or defective mitochondrial signaling in the germline. To test this, we first ruled out the occurrence of embryonic or larval lethality in *gsr-1a trxr-2* double mutants (Fig 2e). Next, we measured total brood size and found that *gsr-1a* mutants had reduced progeny as compared to that of wild-type or *trxr-2* mutant controls while *gsr-1a trxr-2* double mutants had the lowest brood size, although it did not differ significantly from that of *gsr-1a* single mutants (Fig. 2f). Finally, due to the delayed time from last molt to egg-lay of *gsr-1a trxr-2* double mutants, we measured the interval during which the four strains produce progeny and found that double mutants have an extended egg-lay period (Fig 2g). Thus, *gsr-1a trxr-2* double mutants show normal somatic development but exhibit delayed reproductive maturation, reduced progeny output and an extended egg-lay window, revealing a specific sensitivity of reproductive physiology associated to mitochondrial redox defects.

**Figure 2.**
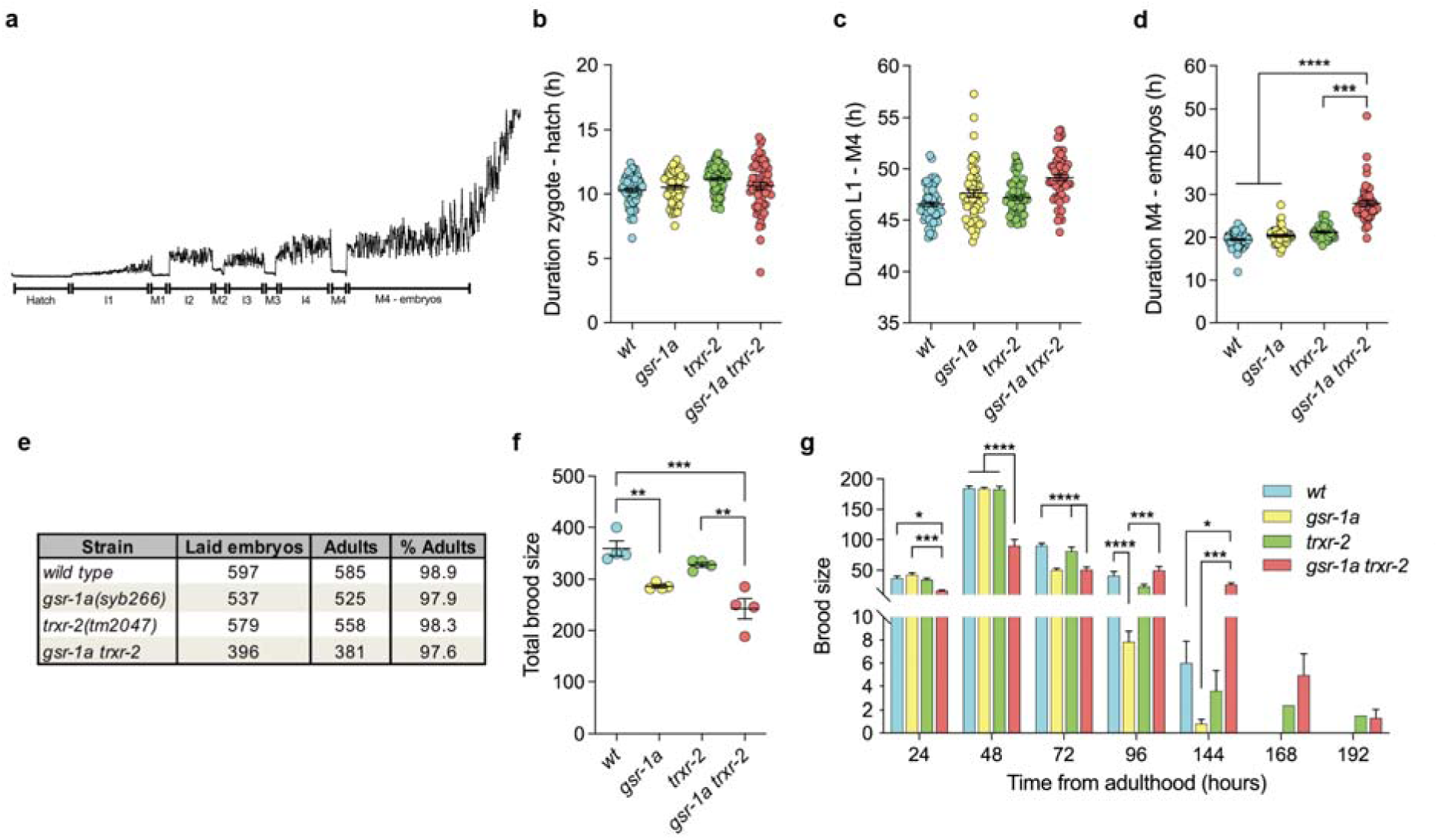
Developmental time and progeny phenotypes of *gsr-1a(syb266)* and *trxr-2(tm2047)* mutants. a) Representative bioluminescence profile of a worm from the zygote stage until the egg-lay period initiation (distorted signal). I1-I4 intermolt intervals, M1-M4 molt intervals, devoid of signal due to halted food ingestion. Quantitative analysis of b) ex-uterus embryonic development, c) larval development, d) egg-lay initiation period. Data are from three different experiments with three biological replicates per assay. Error bars are SEM. *** p<0.001; **** p<0.0001 by ordinary One-way ANOVA (b,c) and by Kruskal-Wallis test (d). e) Percentage of animals that reach adulthood. Data are pooled from two independent experiments with three biological replicates. f) Quantification of the total brood size of 20 animals. Each point represents the average progeny of 5 animals on the same plate. Error bars are SEM. ** p<0.01; *** p<0.001 by ordinary One-way ANOVA. g) Bars show the progeny per day from 20 animals of each genotype. Data are the mean +/− SEM. * p<0.05; *** p<0.001; **** p<0.0001 by ordinary One-way ANOVA.

### Disruption of mitochondrial redox homeostasis induces mitochondrial UPR

To gain deeper insight at the molecular level on the consequences of disrupting redox homeostasis in mitochondria, we performed a RNAseq analysis. Consistent with the lack of phenotype in *gsr-1a* and *trxr-2* single mutants, we only observed a minor alteration of the mRNA expression pattern in *gsr-1a* or *trxr-2* animals (Fig 3a). In contrast, *gsr-1a trxr-2* double mutants showed a dramatic alteration of their transcriptional profile with hundreds of genes up and downregulated, some of which were validated by qPCR (Fig 3, Sup Fig 2a,b and Supplementary Tables 2,3). To identify the biological processes and the biochemical pathways in which these genes participate we performed a Gene Ontology and KEGG enrichment analysis (Gene Ontology et al. 2023; Kanehisa et al. 2025) and found a strong upregulation of genes related to protein phosphorylation, detoxification and immune stress response, transmembrane receptor signaling, male gamete generation, metabolism of sulfur containing amino acids and cuticle development. In turn, downregulated genes were mainly involved in proteasomal function and cell specification processes (Fig 3b).

**Figure 3.**
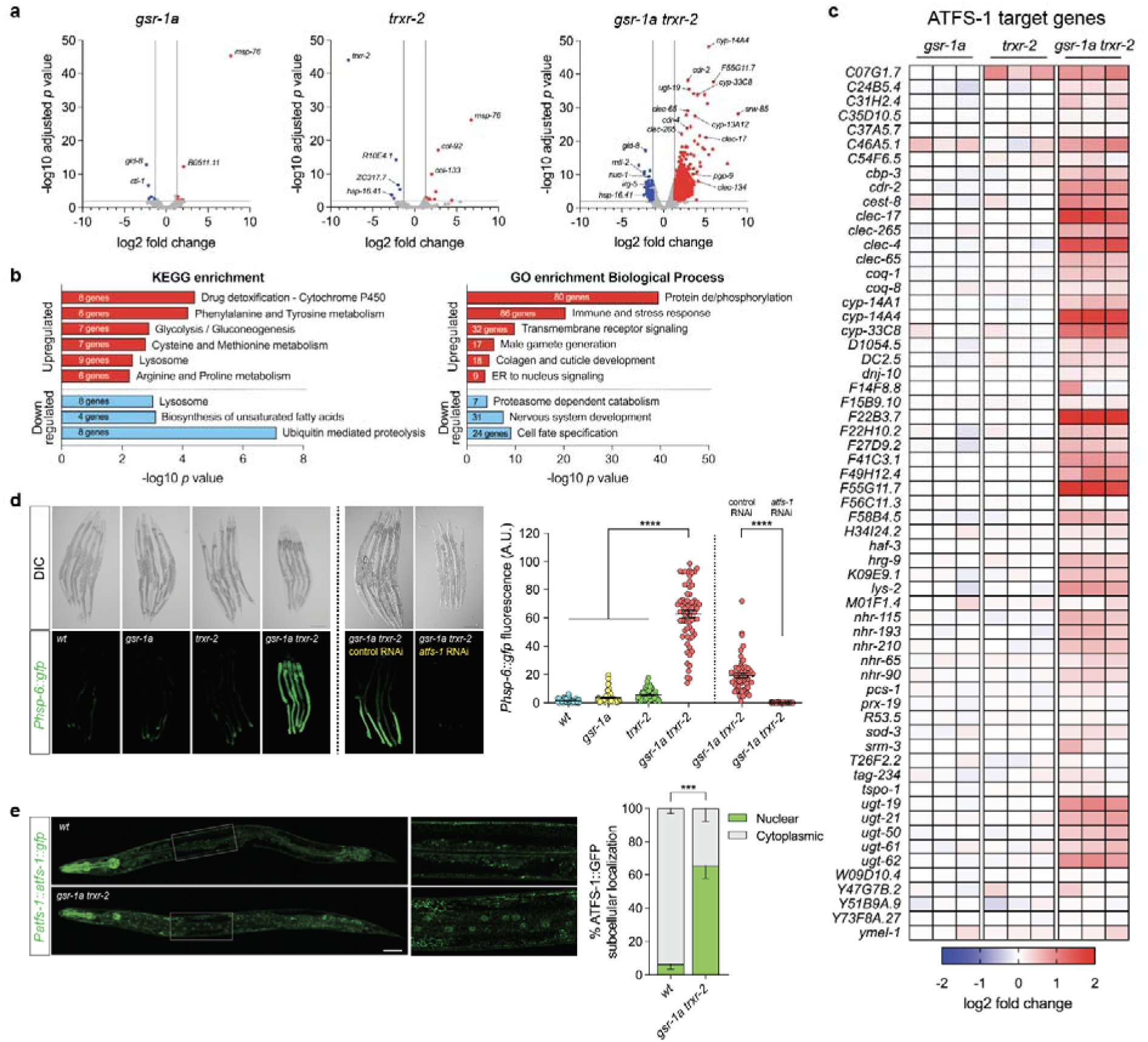
Transcriptomic analysis of *gsr-1a(syb266)* and *trxr-2(tm2047)* mutants. a) Volcano-plots of *gsr-1a* and *trxr-2* single and double mutants. The Log2 fold change indicates the mean expression levels for each gene, represented by dots). No differentially expressed genes (DEG) are indicated in grey, upregulated genes in red and downregulated genes in blue. Data are from three independent experiments with three biological replicates analysed with the edgeR package (Robinson et al. 2010). b) Gene Ontology and KEGG pathway enrichment clustering of DEGs (FC>2, p<0.05) between *gsr-1a trxr-2* double mutants and wild-type controls. c) Heat-map of the high confidence ATFS-1 target genes (Soo and Van Raamsdonk 2021). The color-coded heat map represents gene expression differences (log2 fold change) relative to the levels found in wild-type controls. d) Representative differential interference contrast and fluorescence micrographs and quantification of worms at first day of adulthood expressing GFP under the control of the *hsp-6* promoter. Data are from three different experiments with at least 20 animals per assay. Error bars are SEM. **** p<0.0001 by Kruskal-Wallis test. Scale bar 200 µm. e) Representative fluorescence micrographs and quantification of L4 worms expressing ATFS-1::GFP under the control of the *atfs-1* promoter. Data are the mean +/− SEM of three independent experiments with at least 70 animals per genotype. *** p<0.001 by two-tailed Mann-Whitney test. Scale bar 50 µm.

The induction of genes involved in drug detoxification, pathogen infection and mitochondrial repair is a common response to mitochondrial dysfunction as a result of disrupting core cellular functions (Melo and Ruvkun 2012; Liu et al. 2014). Interestingly, among the most upregulated genes in the *gsr-1a trxr-2* double mutant we found many high confidence ATFS-1 target genes (Soo and Van Raamsdonk 2021) (Fig 3c), suggesting that these animals may have an induced UPR^mt^. To test this, we used the UPR^mt^ reporter transgene *zcIs13[Phsp-6::gfp]* (Yoneda et al. 2004) and found that only the *gsr-1a trxr-2* double mutant but not the respective single mutant controls strongly induced GFP expression (Fig 3d). Importantly, this response was fully dependent on ATFS-1 transcription factor as *atfs-1* RNAi downregulation completely abolished the fluorescence in *gsr-1a trxr-2; zcIs13[Phsp-6::gfp]* animals (Fig 3d). Consistently, *gsr-1a trxr-2* double mutants have a constitutive nuclear localization of ATFS-1, prerequisite to trigger a UPR^mt^ transcriptional response (Fig 3e) (Nargund et al. 2012). The relevance of ATFS-1 function in the survival of *gsr-1a trxr-2* worms is highlighted by the fact that a triple mutant *gsr-1a trxr-2; atfs-1* is not viable and can only be maintained as a balanced strain (strain VZ1063 Supplementary Table 1). Moreover, while canonical ATFS-1 dependent UPR^mt^ is strongly induced in *gsr-1a trxr-2* double mutants (approx. 50-fold induction) (Fig. 3d), other mitochondrial related stress responses such as the ATFS-1 independent UPR^mt^ (Munkacsy et al. 2016), mitochondrial to cytosolic stress response (MCSR) (Kim et al. 2016) or ethanol and stress response element (ESRE) (Tjahjono et al. 2020), as well as UPR^er^ were only very marginally altered (Sup Fig 2c).

Collectively, the phenotypic and transcriptomic analyses show that GSR-1a and TRXR-2 are functionally redundant in mitochondria and that their simultaneous absence induces a robust ATFS-1 dependent UPR^mt^ response that is essential for the survival of animals with compromised redox homeostasis in this organelle.

**Supplemental Figure 2.**
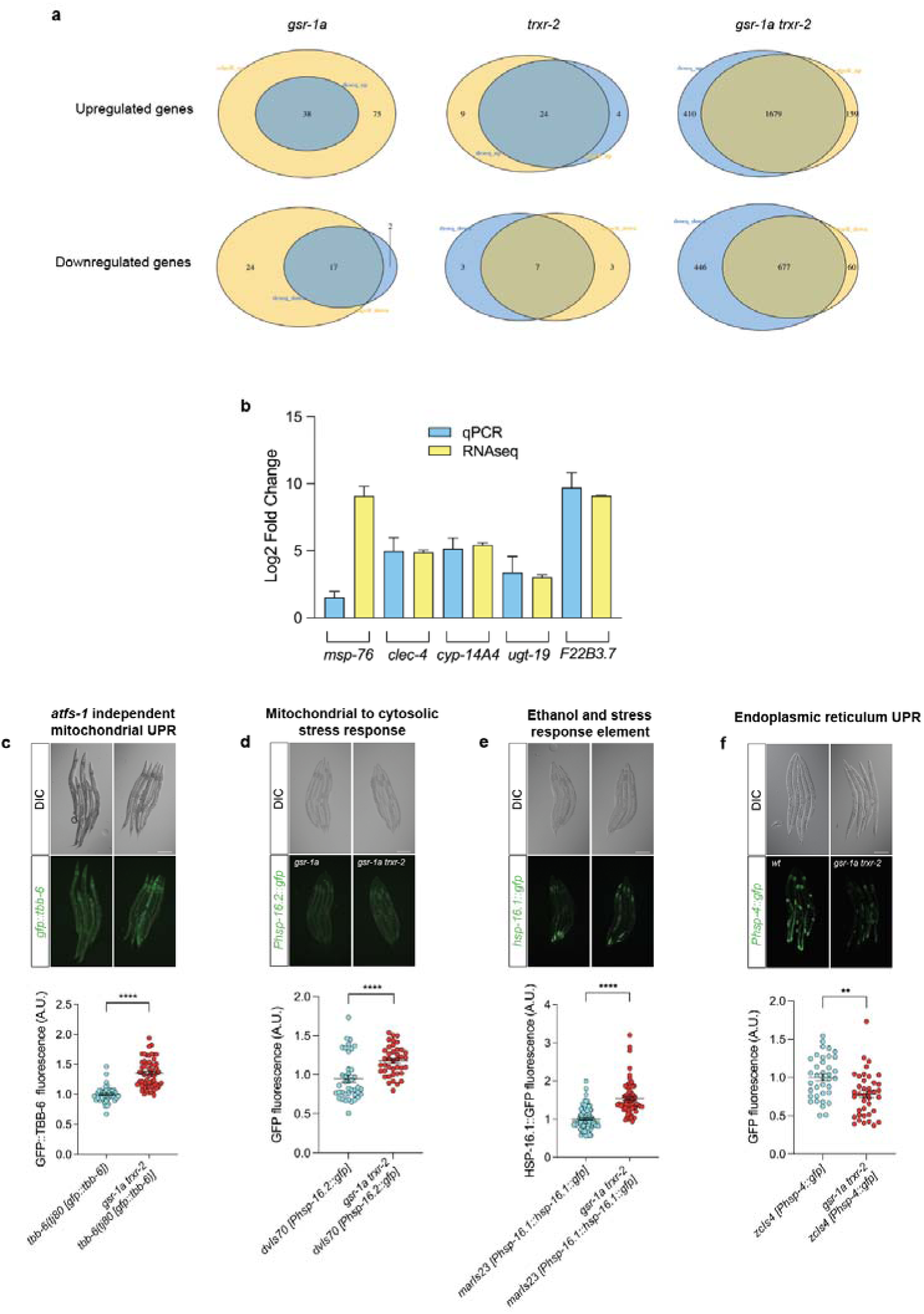
Transcriptomic analysis and stress response markers in *gsr-1a(syb266)* and *trxr-2(tm2047)* mutants. a) Venn diagrams and overlap of DEGs analysed by edgeR and Deseq2 packages (Robinson et al. 2010; Love et al. 2014). b) Comparison of selected DEGs by RNAseq and qPCR analysis. c-f) Representative differential interference contrast and fluorescence micrographs and quantification of worms at first day of adulthood, except *dvIs70* animals at L4 stage, expressing reporters for c) *atfs-1* independent UPR^mt^ (Munkacsy et al. 2016); d) mitochondrial-to-cytosolic stress response (Kim et al. 2016); e) ethanol and stress response element (Tjahjono et al. 2020) and f) UPR^er^ (Yoneda et al. 2004). Data are from three different experiments with at least 10 animals per assay. Error bars are SEM. ** p<0.01; **** p<0.0001 by unpaired t-test. Scale bar 200 µm.

### Induction of mitochondrial UPR does not enhance stress resistance or increase lifespan in *gsr-1a trxr-2* mutants

*C. elegans* continuously monitors the correct function of core cellular activities such as protein synthesis, proteasome-dependent degradation or mitochondrial respiration. Disruption of these essential cellular activities is perceived as an external insult, triggering a defensive transcriptional program that includes genes involved in pathogen resistance, xenobiotic detoxification and innate immune response (Melo and Ruvkun 2012). The transcriptomic profile of *gsr-1a trxr-2* mutants, which is enriched for genes within these pathways (Fig 3b), resembles the transcriptional responses of mutants compromising core mitochondrial functions such as the electron transport chain, tricarboxylic acid cycle or iron-sulfur cluster biogenesis (Melo and Ruvkun 2012; Liu et al. 2014; Kim et al. 2023; Schiavi et al. 2023). These similarities suggest that maintenance of mitochondrial redox homeostasis is among the key mitochondrial functions under cellular surveillance that are essential for organismal survival.

Because robust transcriptional remodeling and UPR^mt^ activation were observed only found in the *gsr-1a trxr-2* double mutant, but not in the respective single mutants, subsequent analyses focused only on the double mutant. We first assessed mitochondrial content and found it slightly decreased in the *gsr-1a trxr-2* worms compared to wild-type controls (Fig 4a). Notably, mitochondrial membrane potential was significantly reduced in the *gsr-1a trxr-2* mutant (Fig 4b), consistent with the UPR^mt^ activation observed in these animals (Berry et al. 2021), whereas mitochondrial ROS production, respiration and ATP levels remained unchanged (Fig 4c-e). Despite the transcriptional activation of stress and defense-related genes, *gsr-1a trxr-2* animals did not exhibit enhanced resistance to the bacterial pathogens *Pseudomonas aeruginosa* or *Staphylococcus aureus* (Figure 4f and Supplementary Table 4), nor to mitochondrial toxicants such as paraquat and juglone (Figure 4g) or the mitochondrial complex I and III inhibitors rotenone and antimycin A, respectively (Figure 4h).

**Figure 4.**
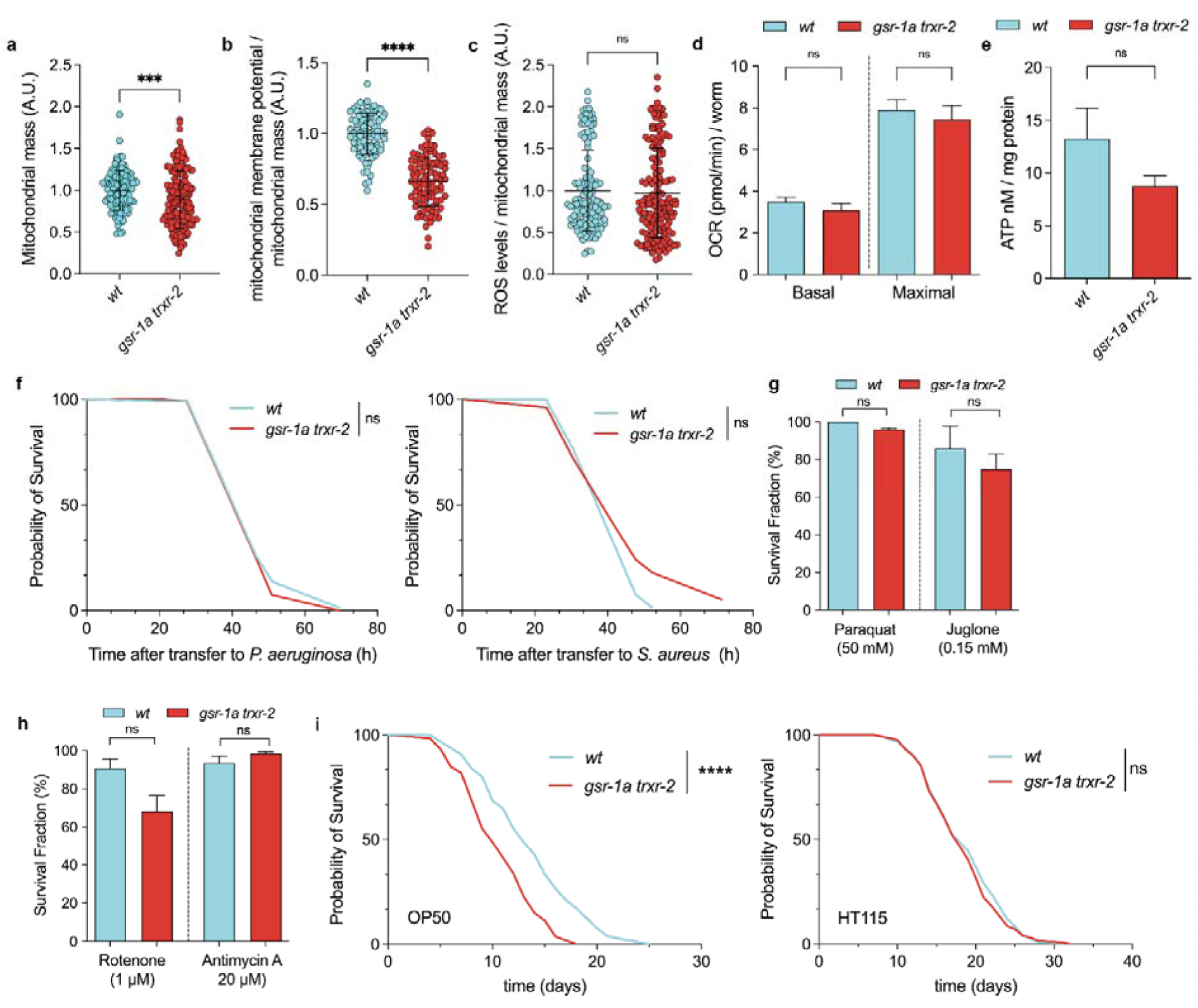
Mitochondrial function, stress response, and lifespan in *gsr-1a(syb266) trxr-2(tm2047)* mutants. a) Quantification of mitochondrial mass using MitoTracker™ Green, b) mitochondrial membrane potential quantification using TMRE and c) mitochondrial ROS levels using MitoTracker™ Red CM-H_2_XRos. All measurements were performed on first day adults and data are from three different experiments with at least 40 animals per assay. ns, not significant; ***p<0.001 and ****p<0.0001 by two-tailed Mann–Whitney test. Error bars are SEM. d) Basal/maximal oxygen consumption rate normalized to the number of worms and e) ATP content normalized by total protein amount. Data are the mean +/− SEM from three different experiments with at least 50 animals per assay. ns, not significant by ordinary One-way ANOVA (d) or two-tailed Mann–Whitney test (e). f) Survival curves following exposure to *Pseudomonas aeruginosa* (left) and *Staphylococcus aureus* (right). Data are from three independent assays for *P. aeruginosa* and two for *S. aureus*, respectively. ns, not significant by Log-rank (Mantel–Cox) test. g) Quantification of survival after 16 hours exposure to paraquat and juglone and h) to rotenone and antimycin. All measurements were performed on L4 animals and data are the mean +/− SEM from three different experiments with at least 40 animals per assay. ns, not significant and *p<0.05 by two-tailed Mann–Whitney test. i) Survival curves of animals fed on *E. coli* OP50 (left) or HT115 (right) bacteria. Data are from three independent assays on OP50 and two on HT115. ns, not significant and ****p < 0.0001 by Log-rank (Mantel–Cox) test with Bonferroni correction.

The relationship between UPR^mt^ induction, typically through depletion of mitochondrial genes, and lifespan modulation is well established, with outcomes depending on the degree of mitochondrial perturbation (Rea et al. 2007; Runkel et al. 2014; Bennett and Kaeberlein 2014). Worms with impaired mitochondrial function often display reduced body size and reduced progeny yet increased lifespan (reviewed in (Haynes and Hekimi 2022)). We therefore tested whether disruption of mitochondrial redox homeostasis also affects longevity. Unexpectedly, we found that *gsr-1a trxr-2* worms are short lived compared to wild-type animals when grown on OP50 bacteria, whereas no differences were detected on HT115 bacteria (Fig 4i Supplementary Table 5). These results suggest that the impact of impairing mitochondrial redox homeostasis on lifespan may be modulated by nutritional cues.

Taken together, these findings indicate that loss of mitochondrial redox homeostasis does not markedly affect stress resistance or mitochondrial physiology in *C. elegans*, likely due to compensatory activation of UPR^mt^-dependent protective pathways, and consistent with the lethality of the *gsr-1a trxr-2; atfs-1* triple mutant.

### Impairment of mitochondrial redox homeostasis alters mitochondrial dynamics without disturbing mitophagy

The observation that *gsr-1a trxr-2* animals are not sensitized to stress despite activation of UPR^mt^ prompted us to investigate whether changes in mitochondrial dynamics, which can be regulated by redox mechanisms (Willems et al. 2015), underlie the preservation of mitochondrial function in these mutants. To this end, we focused on muscle and hypodermal cells, two tissues with high metabolic activity and energy demand, as muscle contractility is required for animal continuous movement, whereas the hypodermis plays a central role in stress resistance, coordinating systemic responses with other tissues through the release of metabolic and hormonal signals (Wang et al. 2024a; Gieseler et al. 2017; Zhang et al. 2023).

Interestingly, *gsr-1a trxr-2* double mutants exhibited tissue-specific alterations in mitochondrial morphology with muscle cells having more elongated mitochondria as compared to wild-type controls, whereas those in hypodermal cells displayed increased fragmentation (Figure 5a,b). This differential morphology was confirmed by quantifying mitochondrial circularity, a widely used parameter for assessing mitochondrial network integrity (Figure 5c) (Teoh et al. 2019). Mitochondrial dynamics are governed by the balance between mitochondrial biogenesis and degradation (mitophagy), as well as by the rates of fusion and fission events, processes modulated by the UPR^mt^ (Naresh and Haynes 2019; Traa et al. 2024; Campbell and Zuryn 2024). In this context, disruption of mitochondrial fission using *drp-1* mutants or mitochondrial fusion with *fzo-1* mutants severely compromised the development of *gsr-1a trxr-2* double mutants (Figure 5d,e) whereas inhibition of mitophagy through *dct-1* or *pink-1* mutations had no effect (Figure 5f,g). Indeed, mitophagy is only slightly increased in *gsr-1a trxr-2* double mutants as shown by the ratio between pH sensitive GFP (autophagosome) and pH-insensitive DsRed (autolysosome) markers of the mtRosella reporter (Sup Fig 3) (Palikaras et al. 2015).

**Figure 5.**
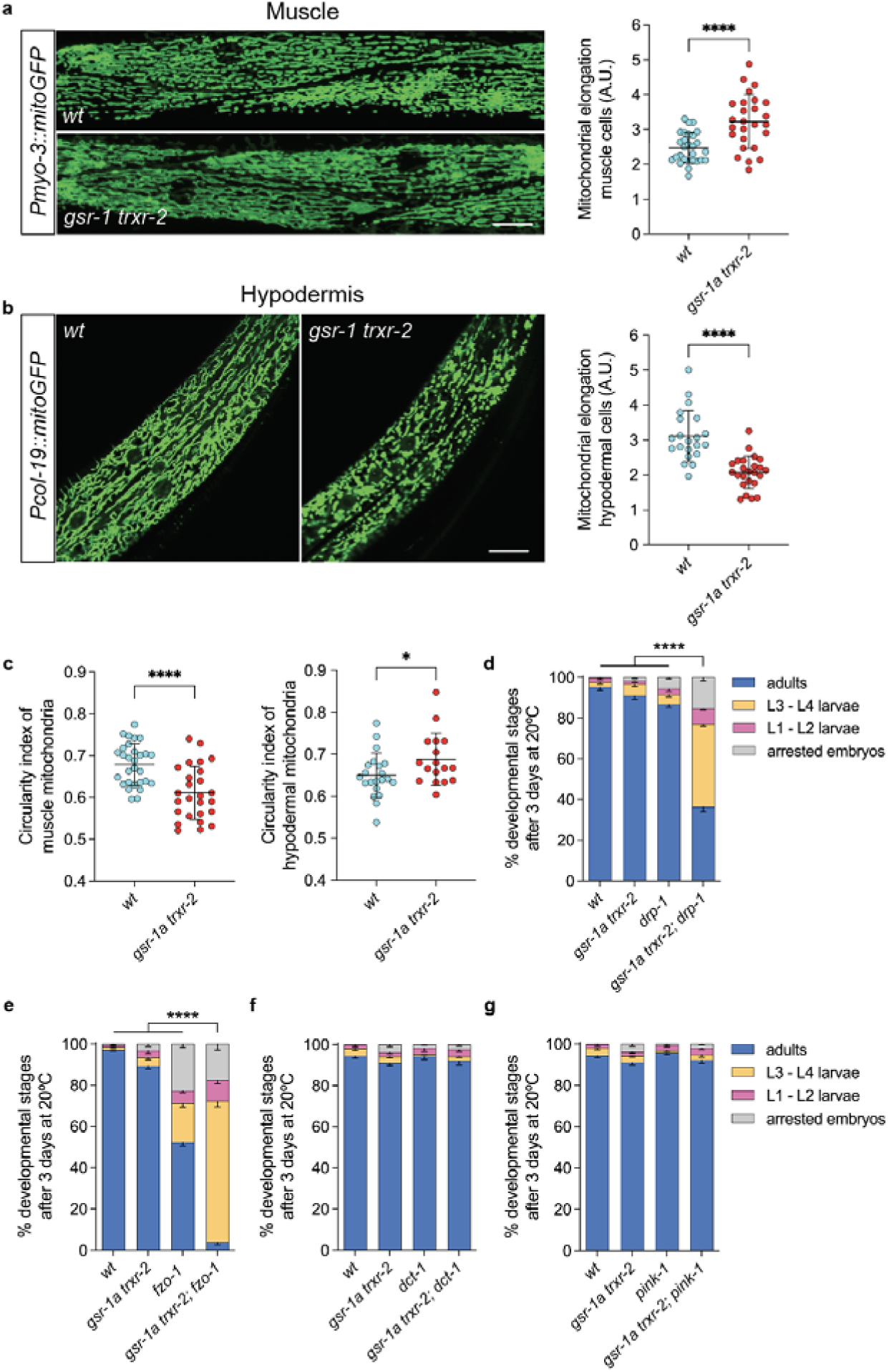
Mitochondrial tissue-specific dynamics of *gsr-1a(syb266) trxr-2(tm2047)* mutants. Representative fluorescence micrographs of mitochondrial network in a) body wall muscle cells and b) hypodermal cells. Graphs show the quantification of mitochondrial elongation from at least 20 first day adults for both tissues. ****p<0.0001 by unpaired t-test. Error bars are SEM. Scale bar 20 µm. c) Quantification of the circularity index of mitochondrial from muscle (left) and hypodermal (right) cells. Data are from at least 20 first day adults. *p<0.05; ****p<0.0001 by unpaired t-test. Error bars are SEM. d-g) Percentage of developmental stages in worms with defective mitochondrial fission-fusion and mitophagy. Data are the mean +/− SEM from three different experiments with at least 100 animals per assay. ****p<0.0001 by ordinary One-way ANOVA.

Together, these findings indicate that mitochondrial network dynamics is tightly regulated in a tissue-specific manner upon disruption of mitochondrial redox homeostasis, even within tissues that are anatomically associated and functionally coordinated such as muscle and hypodermis (Francis and Waterston 1991). Moreover, the dispensability of mitophagy for *gsr-1a trxr-2* double mutants development supports the notion that overall mitochondrial function remains largely preserved, precluding the need for selective degradation of damaged mitochondria, or alternatively, that compensatory mechanisms such as exopher-mediated mitochondria extrusion (Melentijevic et al. 2017) contribute to the maintenance of a healthy mitochondrial network.

**Figure 3S.**
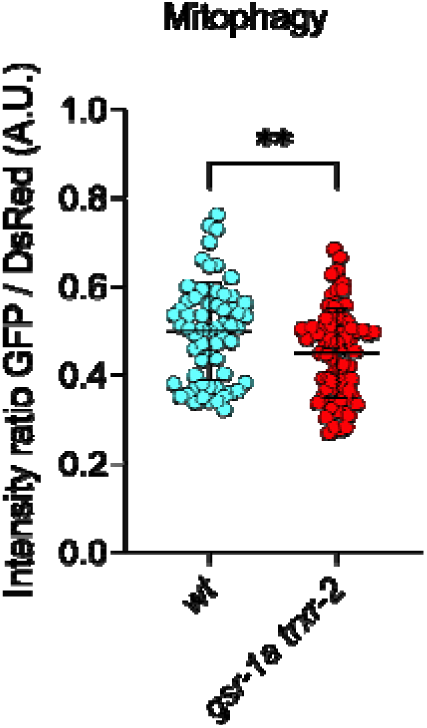
Quantification of mtRosella biosensor. Data are from three independent experiments with at least 10 first day adult animals per assay. Represented is the ratio between pH-sensitive GFP to pH-insensitive DsRed. **p<0.01 by unpaired t-test. Error bars are SEM.

### Tissue-specific somatic and germline effects upon impairment of mitochondrial redox homeostasis

Given the differential effects of the *gsr-1a trxr-2* deficiency on mitochondrial dynamics in muscle and hypodermal, we next focused on phenotypes specifically associated to these two tissues. Using the CeleST software, which quantifies locomotory behaviour in liquid medium as a proxy for muscle function (Restif et al. 2014), we observed that first day adult *gsr-1a trxr-2* animals exhibit reduced locomotor fitness parameters together with an increase in muscle frailty indicators (Figure 6a,b). These findings suggest a functional decline of muscle cells when mitochondrial redox homeostasis is compromised. Consistent with this interpretation, *gsr-1a trxr-2* double mutants also display a reduced pharyngeal pumping rate (Figure 6c), a defect that is not attributable to altered muscle depolarization capacity by defective neuronal signaling, as demonstrated by electropharyngeogram measurements (Figure 6d and Sup Fig 4a).

**Figure 6.**
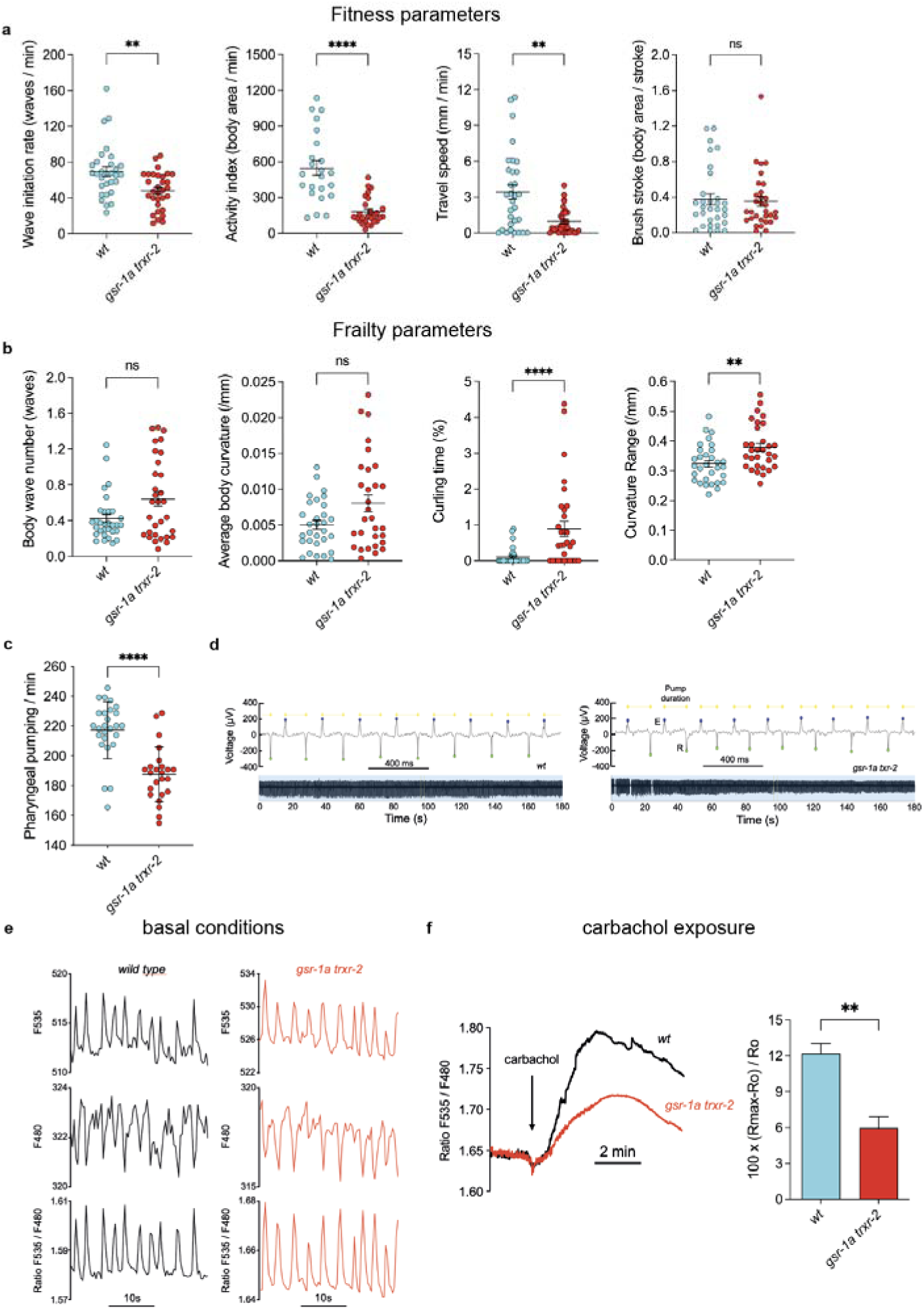
Muscle-specific phenotypes in *gsr-1a(syb266) trxr-2(tm2047)* mutants. CeleST locomotory parameters associated with a) physiological fitness and b) physiological frailty in first day adult animals. Data are from three independent experiments with at least 10 animals per assay. ns, not significant; **p<0.01; ****p<0.0001 by two-tailed Mann-Whitney test. Error bars are SEM. c) Quantification of pharyngeal pumping rate per minute. Data are from three independent experiments with at least 10 animals per assay. ****p<0.0001 by two-tailed Mann-Whitney test. Error bars are SEM. d) Representative electropharyngeograms (EPGs) measurements performed in animals at fourth day of adulthood. Measurements were performed on 31 wild-type worms and 30 *gsr-1a trxr-2* double mutants. The following parameters were obtained and pooled: frequency, pump duration, amplitude (voltage change from the E wave peak to the R wave peak) and the R/E ratio. The mean data for these parameters are shown in Fig. 4S. e) Ca^2+^ oscillations under resting conditions in pharyngeal muscle cells. The figure shows typical records of the two individual F535 and F480 fluorescences (in arbitrary units), and the F535/F480 ratio, which directly reflects changes in Ca^2+^ concentration. Measurements were performed on 24 wild-type worms and 32 *gsr-1a trxr-2* double mutants. The following parameters were obtained and pooled: frequency, width at baseline, width at half-height and height. The mean data for these parameters are shown in Fig. 4Sb. f) Effect of carbachol on mitochondrial Ca^2+^ uptake. The figure shows the average of 17 experiments performed on wild-type worms and 20 experiments performed on *gsr-1a trxr-2* double mutants. Carbachol was added to the chamber when indicated to reach a final concentration of 10mM. The bar diagram illustrates the height of the Ca^2+^ peak induced by carbachol in both conditions. **p<0.001 by unpaired t-test.

Proper regulation of mitochondrial calcium uptake is essential for muscle performance, both to sustain ATP production during contraction and to maintain cytosolic calcium homeostasis (Montero et al. 2000; Garbincius and Elrod 2022). Using a mitochondrial-targeted fluorescence reporter expressed in pharyngeal muscle cells (Alvarez-Illera et al. 2017), we found that *gsr-1a trxr-2* animals show normal basal mitochondrial calcium import compared to wild-type controls (Figure 6e and Sup Fig 4b). However, when worms are exposed to carbachol, a chemical that induces a massive calcium entry into the cytoplasm by activation of the nicotinic acetylcholine receptors in the pharyngeal muscle cells (Raizen et al. 1995), we observed a marked reduction in mitochondrial calcium uptake in the *gsr-1a trxr-2* double mutants (Figure 6f). Together, these findings indicate that mitochondrial redox homeostasis is required to efficiently buffering high levels of calcium in the cytoplasm.

**Figure 4S:**
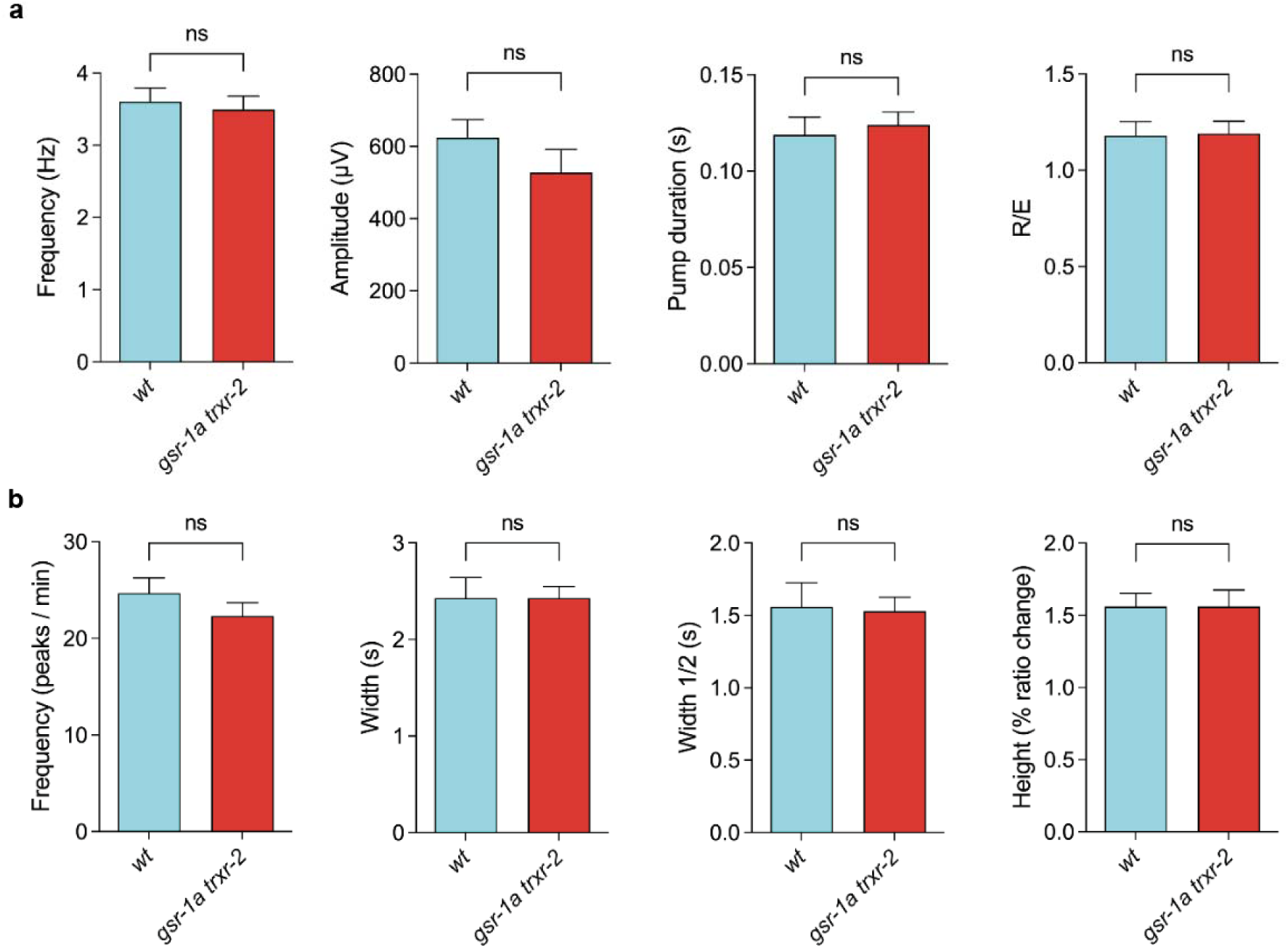
Electropharyngeogram and basal mitochondrial calcium uptake parameters in wild type and *gsr-1a trxr-2* mutants. a) Quantification of electropharyngeogram frequency, pump duration, amplitude and repolarization (R) / depolarization (E) ratio. Data are the mean +/− SEM from experiments performed in 31 wild-type worms and 30 *gsr-1a trxr-2* double mutants. ns, not significant by unpaired t-test. b) Quantification of basal mitochondrial Ca^2+^ peaks: frequency, width at baseline (width), width at half-height (width ½) and height. Data are the mean +/− SEM from experiments performed in 24 wild-type worms and 32 *gsr-1a trxr-2* double mutants. ns, not significant by unpaired t-test.

Reduced body size is a well characterized phenotype associated with impaired mitochondrial function (Dillin et al. 2002), and this trait is also observed in *gsr-1a trxr-2* double mutants. In *C. elegans*, body size is primarily regulated by the BMP/DBL-1 signaling pathway, which promotes endoreduplication of hypodermal cells and transcriptional control of collagen genes, the major structural components of worm cuticle, synthesized by the underlying hypodermis (Madaan et al. 2018; Page and Johnstone 2007; Flemming et al. 2000). Interestingly, our RNAseq analysis revealed upregulation of genes encoding collagens and other proteins involved in cuticle morphogenesis (Figure 3b and Supplementary Tables 2,3). Notably, several of these genes, including *lon-3* and *sqt-1*, function downstream BMP/DBL-1 pathway to control worm body size (Suzuki et al. 2002; Kramer et al. 1988). As elevated *lon-3* expression is known to decrease body size (Nystrom et al. 2002), we hypothesized that increased *lon-3* activity contributes to the reduced size phenotype observed in *gsr-1a trxr-2* animals. To test this, we used the *lon-3(sp23)* loss-of-function mutation (Nystrom et al. 2002) and found that the increased ratio in body size of the *gsr-1a trxr-2; lon-3* triple mutant compared to *gsr-1a trxr-1* control was similar to that of *lon-3* single mutant compared to wild-type control (Figure 7a), indicating that *lon-3* is not responsible for the small phenotype. Furthermore, *gsr-1a trxr-2* double mutants have the same number of nuclei in the hypodermal hyp-7 syncytium as wild-type animals (Figure 7b), suggesting that neither defective cuticle formation nor impaired hypodermal DNA content (ploidy) accounts for the reduced body size of *gsr-1a trxr-2* worms. Given the high number of collagen encoding genes present in the *C. elegans* genome (Page and Johnstone 2007), we cannot rule out that redox regulation of collagens other than *lon-3* may control body size in the double mutants.

**Figure 7.**
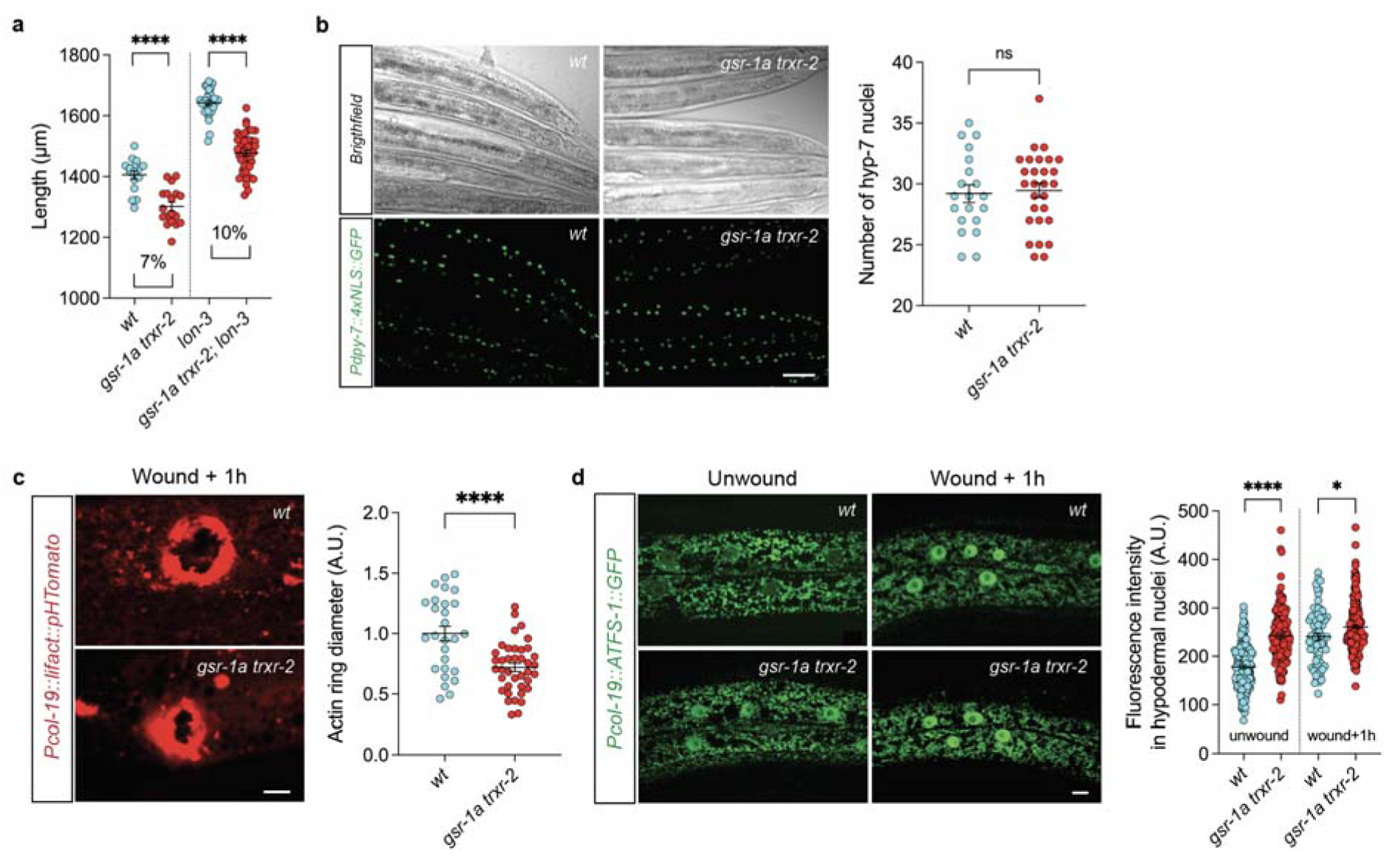
Hypodermal-dependent phenotypes of *gsr-1a(syb266) trxr-2(tm2047)* mutants. a) Quantification of day 2 adult animals length. Percentages represent the differential length between indicated genotypes. Data are from three independent experiments with at least 15 animals per assay. ****p<0.0001 by two-tailed Mann-Whitney test. Error bars are SEM. b) Representative bright-field and fluorescence micrographs (left) and quantification of syncytium nuclei (right) of the posterior part of hyp-7 cell. Data are from at least 20 animals. ns, not significant by unpaired t-test. Error bars are SEM. Scale bar 25 µm. c) Representative fluorescence micrographs (left) and quantification of the diameter of the laser induced wound after 1 hour recovery (right). Data are from at least 25 first day adult animals. ****p<0.0001 by unpaired t-test. Error bars are SEM. Scale bar 10 µm. d) Representative fluorescence micrographs (left) and quantification of the fluorescence intensity of the ATFS-1::GFP reporter in hypodermal cell nuclei before and after 1 hour laser induced wound. Data are from at least 65 first day adult animals. *p<0.05; ****p<0.0001 by unpaired t-test. Error bars are SEM. Scale bar 20 µm.

Hypodermal wounding induces a distinct innate immune response initiated by a mitochondrial ROS production at the site of injury, triggering a rapid and reversible mitochondrial fragmentation that depends on cytoplasmic calcium and the mitochondrial Rho GTPase MIRO-1 (Pujol et al. 2008; Xu and Chisholm 2014a; Fu et al. 2020). As *gsr-1a trxr-2* double mutants induce an immune response at the transcriptional level and also show increased mitochondrial fragmentation in hypodermis, we next asked whether wound repair dynamics is altered in these animals. Indeed, laser-wounding experiments revealed that *gsr-1a trxr-2* worms repair wounds more rapidly than wild-type controls, as measured by contraction of the actin right at the injury site (Figure 7c). Additionally, wounding triggered nuclear translocation of an ATFS-1::GFP fusion protein in hypodermal cells, a phenotype that is also found in unwounded *gsr-1a trxr-2* double mutants (Figure 7d). Together, these observations suggest that accelerated wound repair in *gsr-1a trxr-2* mutants may be driven by chronic activation of ATFS-1-dependent stress signaling in the hypodermis.

*gsr-1a trxr-2* double mutants have decreased progeny production, a phenotype previously associated with impaired mitochondrial function in *C. elegans* (Curran et al. 2004). Because our RNAseq analysis revealed an upregulation of genes involved in male gamete generation and function (Figure 3b), we first examined sperm production and fertilization capacity in *gsr-1a trxr-2* animals. Although *gsr-1a trxr-2* double mutants produced a comparable number of spermatozoa as compared to wild-type controls, their fertilization efficiency was markedly reduced (Figure 8a,b). To evaluate whether maternal mechanisms also account to the lower progeny of *gsr-1a trxr-2* double mutants we next quantified exopher production in these animals. As described previously, exophers represent a specialised class of exceptionally large extracellular vesicles that enable *C. elegans* cells to externalise material that cannot be efficiently processed by conventional intracellular quality-control pathways. First described in neurons as a mechanism for expelling damaged mitochondria and protein aggregates (Melentijevic et al. 2017), exophers are also produced by body-wall muscle, where they serve dual functions: the removal of mitochondria and the export of yolk proteins to support embryonic development (Turek et al. 2021). As shown in Figure 8c, *gsr-1a trxr-2* double mutants generate fewer muscle-derived exophers compared to wild-type controls. Given the induction of UPR^mt^ in *gsr-1a trxr-2* animals, we next examined whether constitutive activation of ATFS-1 influences exopher formation. In *atfs-1(et17)* animals, which express an ATFS-1 variant with a defective mitochondrial targeting sequence leading to constitutive ATFS-1 nuclear localization, the number of muscle exopher formation was also reduced (Figure 8d). This finding provides a potential mechanistic link between disrupted mitochondrial redox homeostasis and impaired exopher formation that may reduce yolk proteins provision to the developing embryos, perhaps by shifting reproductive resources to mitochondrial repair. Taken together, our data suggest that the diminished progeny production in *gsr-1a trxr-2* animals can result from both reduced sperm fertilization capacity and decreased muscle-derive exopher formation

**Figure 8.**
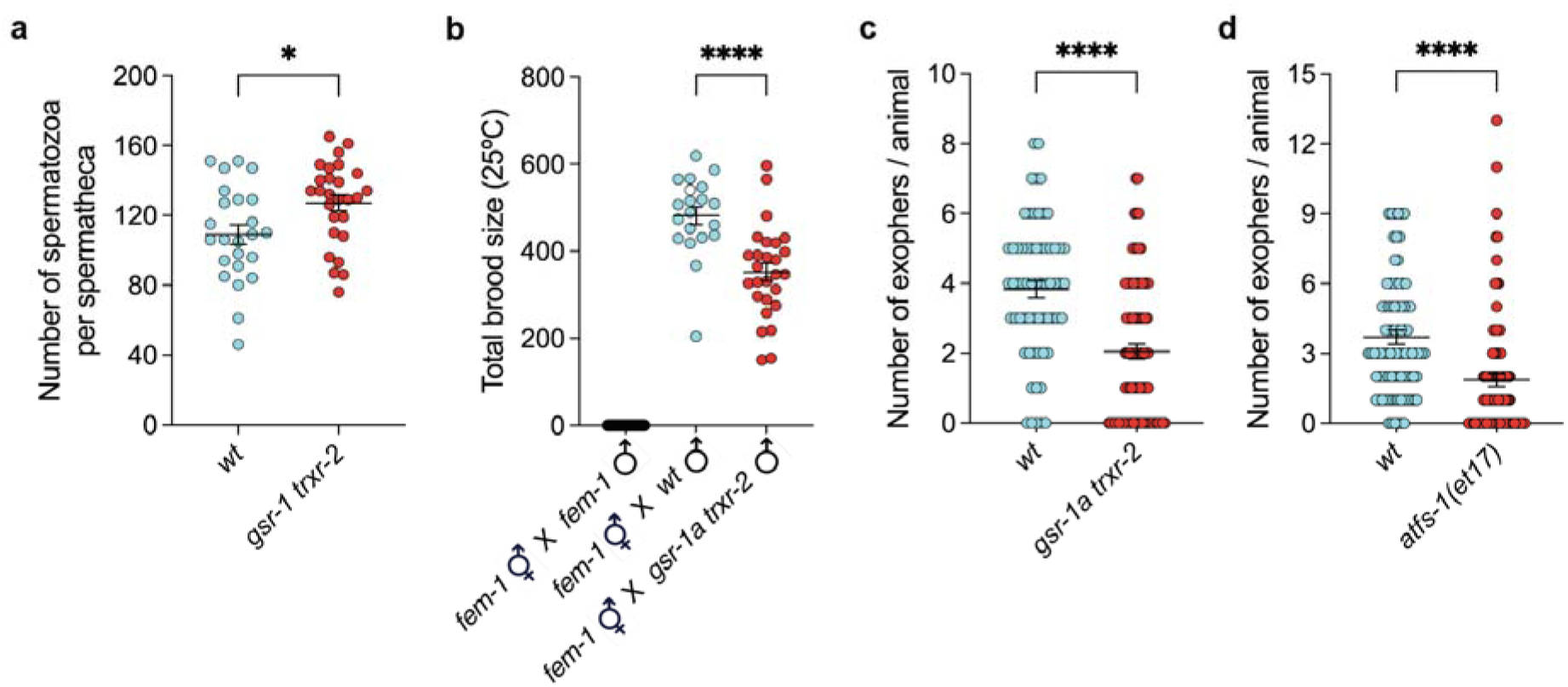
Fertilization capacity and exopher production phenotypes in *gsr-1a(syb266) trxr-2(tm2047)* mutants. a) Quantification of the number of spermatozoa per spermatheca. Data are from four independent experiments with at least four spermathecas per assay. *p < 0.05 by unpaired t-test. Error bars are SEM. b) Quantification of the total brood size from *fem-1(hz17)* hermaphrodites fertilized at 25 °C. Data are from three independent experiments including at least 10 crosses and progeny production was monitored over eleven days. ****p< 0.0001 by unpaired t-test. Error bars are SEM. c,d) Quantification of muscle exopher formation. Data are from three independent experiments with at least ten animals per assay. ****p< 0.0001 by two-tailed Mann-Whitney test. Error bars are SEM.

In summary, while disruption of mitochondrial redox homeostasis compromises muscle and sperm function, it may simultaneously promote protective adaptations in other tissues, such as hypodermis.

## Discussion

Mitochondria rely on the import of reduced glutathione from the cytoplasm, where it is synthesized. Accordingly, mammalian cells lacking the mitochondrial GSH importer paralogues SLC25A39 and SLC25A40 fail to proliferate (Wang et al. 2021), and this phenotype is conserved in *C. elegans* as null mutants of the *C16C10.1* gene, the single worm orthologue of SLC25A39/40 genes, die at L1 stage. In *C. elegans*, deletion of *gsr-1* first exon selectively abolishes the expression of the mitochondrial GSR-1a isoform while maintaining the expression of the cytoplasmic isoform GSR-1b, which is essential for survival (Mora-Lorca et al. 2016). Interestingly, *gsr-1a* mutants exhibit minimal transcriptional changes, similarly to *trxr-2* mutants lacking mitochondrial thioredoxin reductase. In contrast, the *gsr-1a trxr-2* double mutant has a robust alteration of its transcriptome, indicating that, like in the cytoplasm, these two redox systems act redundantly to maintain mitochondrial redox homeostasis.

Because the primary function of glutathione reductase is to regenerate GSH from GSSG, the observed transcriptional reprogramming in *gsr-1a trxr-2* double mutants suggests that the mitochondrial thioredoxin system may substitute this GSSG recycling function in the absence of mitochondrial GSR-1a. The viability of *gsr-1a trxr-2* animals, with only modest physiological defects, could be explained by the existence of a third GSSG reduction system in mitochondria either directly by dihydrolipoic acid or via mitochondrial glutaredoxin GLRX-5 and dihydrolipoamide (Bast and Haenen 1988; Porras et al. 2002). Alternatively, an active GSSG export mechanism may operate in these animals to reduce it in the cytoplasm. Indeed, *C. elegans* mutants lacking the putative GSSG exporter *abtm-1* are lethal (Gonzalez-Cabo et al. 2011). Together, it is then plausible that mitochondrial thioredoxin reductase and glutathione reductase systems are more involved in general redox homeostasis maintenance rather than recycling mitochondrial GSSG, which can then be performed in the cytoplasm if efficiently exported from mitochondria or within mitochondria through the lipoic acid cycle.

The transcriptional signature of the *gsr-1a trxr-2* double mutant resembles that of other mutants that impair core mitochondrial functions, including genes involved in stress resistance and immune response (Melo and Ruvkun 2012; Liu et al. 2014; Kim et al. 2023; Schiavi et al. 2023). Consistent with this, *gsr-1a trxr-2* animals are smaller and display germline developmental defects that reduce progeny production and extend the egg-laying period, phenotypes associated with impaired mitochondrial function (Rea et al. 2007; Dillin et al. 2002). Related to this, and specific to mitochondrial impairment, mitohormesis is an adaptive response by which a mild, sublethal mitochondrial stress triggers a number of cellular changes leading to a rewiring of cell metabolism, protein folding capacity and redox environment that collectively predispose the organism to be less vulnerable to subsequent more severe stresses (reviewed in (Cheng et al. 2023)). However, despite *gsr-1a trxr-2* double mutants are small and have reduced progeny, phenotypes shared by mitohormetic worms (Yang and Hekimi 2010; Ventura et al. 2005), they do not show enhanced resistance to pathogen infection or toxicants exposure, do not live longer and exhibit normal ROS levels and basal respiration. Thus, disruption of mitochondrial redox homeostasis alters mitochondrial function but does not seem to elicit a beneficial adaptive mitohormetic response.

*gsr-1a trxr-2* animals trigger a transcriptional response consistent with the hypothesis that maintenance of redox homeostasis is a core mitochondrial function, as its impairment triggers the expression of many genes that are *bona-fide* targets of ATFS-1, the master regulator of UPR^mt^ (Soo and Van Raamsdonk 2021). To our knowledge, this genetic background induces one of the strongest constitutive ATFS-1 nuclear import phenotypes described to date. These findings highlight the central role of mitochondrial redox balance in sustaining proteostasis and demonstrate the essential role of ATFS-1 activation and UPR^mt^ induction during development when mitochondrial redox homeostasis is disrupted, proven by the lethal phenotype of a *gsr-1a trxr-2; atfs-1* triple mutant.

Throughout development, *C. elegans* undergoes a mitochondrial network expansion program, particularly critical during germline maturation at the L3 to L4 larval stage transition, which requires ATFS-1 mediated UPR^m^ induction (Charmpilas and Tavernarakis 2020; Shpilka et al. 2021). Our data support the hypothesis that a tightly regulated mitochondrial redox function is essential for a correct germline development and acquisition of normal size and that the constitutive UPR^mt^ induction in *gsr-1a trxr-2* animals can only partly restore mitochondrial functionally to compensate germline and size defects. This partial compensation likely underlies the reduced body size, impaired gonad migration and decreased progeny phenotypes observed in the double mutants, reflecting a trade-off between stress mitigation and reproductive fitness, which is dependent on ATFS-1 and UPR^mt^ (Zhou and Liu 2025).

We further found that mitochondrial dynamics in *gsr-1a trxr-2* double mutants is regulated in a tissue-specific manner, with muscle cells showing increased elongated mitochondria while hypodermal cells presenting a high degree of mitochondrial fragmentation. Proper mitochondrial dynamics is essential for muscle performance, as illustrated by the motility defects described in mutants that either disrupt mitochondrial fusion (*fzo-1*) or mitochondrial fission (*drp-1*) (Byrne et al. 2019). In this context, *gsr-1a trxr-2* double mutants also have impaired motility phenotypes and other muscle related defects such as reduced pharyngeal pumping and, more strikingly, display severe developmental delays when combined with mutations in *drp-1* or *fzo-1* genes, indicating that mitochondrial redox imbalance sensitizes mitochondrial dynamics. In contrast, the small increase of mitophagy in *gsr-1a trxr-2* double mutants does not have any impact on development, demonstrating that disruption of mitochondrial redox homeostasis primarily alters mitochondrial remodeling rather than causing accumulation of damaged mitochondria.

Interestingly, calcium influx into the mitochondrial matrix, which is important to provide energy for muscle contraction, is not altered under basal conditions in *gsr-1a trxr-2* animals, but it is strongly decreased upon carbachol treatment, that induces a massive calcium influx through the plasma membrane and the subsequent buffering by import into mitochondria (Raizen et al. 1995; Montero et al. 2000). Our data suggest that MCU-1 inner mitochondrial membrane Ca^2+^ uniporter, which mediates calcium import into mitochondrial matrix upon carbachol treatment (Alvarez-Illera et al. 2020), is negatively regulated when mitochondrial redox homeostasis is disrupted. The fact that mammalian MCU is activated by glutathionylation in a conserved cysteine residue (Dong et al. 2017), also present in *C. elegans*, may provide a plausible explanation for the lower Ca^2+^ import in *gsr-1a trxr-2* double mutants as MCU-1 glutathionylation may be defective in muscle cells.

In contrast to the increased elongation of the mitochondria network in muscle cells, hypodermal cells have a much more fragmented network in *gsr-1a trxr-2* worms as compared to wild-type animals. Unexpectedly, *gsr-1a trxr-2* animals are more efficient repairing laser-induced lesions in the hypodermis, which may appear as a counterintuitive result considering that closure of hypodermal wounds has been shown to be dependent on Ca^2+^ import into the mitochondria via MCU-1 (Xu and Chisholm 2014a). One possible explanation for this apparently contradictory result compared to muscle cells is that laser-wounding, a much more damaging treatment than carbachol exposure, may also induce an increase in cytoplasmic GSH import into mitochondria to sustain MCU-1 activity, as it has been reported following traumatic brain injury in mice (Wang et al. 2025). Alternatively, enhanced mitochondrial fragmentation in hypodermis may induce MCU-1 levels in this tissue, while mitochondrial elongation may decrease MCU-1 expression in muscle cells. It will be interesting to determine whether ATFS-1 directly regulates cytoskeletal repair genes or modulates calcium dynamics during wound closure.

In addition to their roles in somatic tissues, mitochondria are essential regulators of *C. elegans* spermatogenesis and sperm activation, the transition from immotile spermatids to fully functional, motile spermatozoa (Wang et al. 2024b; Kelleher et al. 2000). Notably, *gsr-1a trxr-2* double mutants exhibit reduced fertilization capacity, even though they upregulate multiple genes required for the formation of functional gametes as shown by transcriptomic analysis. How defects in mitochondrial redox homeostasis specifically impair sperm maturation and/or sperm function, and whether diminished muscle-derived exopher yolk provisioning to developing embryos contributes to the reduced brood-size phenotype in these mutants, remain important open questions that deserve further investigation.

In conclusion, in this work we provide a genetic and functional characterization a *C. elegans* mutant with impaired mitochondrial redox homeostasis, which leads to mild but pleiotropic phenotypes. The availability of this mutant simultaneously lacking mitochondrial thioredoxin and glutathione reductases offers a valuable model to investigate how mitochondrial redox control influences key cellular and organismal functions. Future studies can build on this work aiming to identify the mitochondria-derived signal responsible for constitutive ATFS-1 activation in *gsr-1a trxr-2* mutants or to examine how endoplasmic reticulum dynamics adapt to compromised mitochondrial redox status. Together, this genetic model provides a useful platform for advancing our understanding of in vivo redox regulation in mitochondria.

## Data availability statement

RNAseq raw data are deposited at the Sequence Read Archive of NCBI with the ID number GSE311484.

## Acknowledgements and funding

Some *C. elegans* strains were provided by the CGC, which is funded by NIH Office of Research Infrastructure Programs (P40 OD010440) USA and by the National BioResearch Project, Japan. We thank SunyBiotech (https://www.sunybiotech.com/) for their excellent assistance generating CRISPR-Cas9 edited alleles and the Genomic Platform at CIBIR for their service and support. We thank Chris Link, Nektarios Tavernarakis, Andrew Chisholm, Encarni Lozano, Ulrich Hartl and Simon Tuck for sharing strains.

MVV was supported by a postdoctoral contract from the Conserjería de Universidad, Investigación e Innovación de la Junta de Andalucía, Spain (DOC_01674). PdlCR was supported by a postdoctoral contract from the Juan de la Cierva Program, Ministerio de Ciencia, Innovación y Universidades, Spain (JDC2023-051269-I). AMV lab was supported by Projects PID2021-122311NB-I00 and PID2024-155904NB-I00, JC lab was supported by Project PID2021-127388NB-I00 and JA lab was supported by Project PID2021-122239OB-I00 (financed by the Spanish Ministerio de Ciencia, Innovación y Universidades (MCIU)), the Spanish Agencia Estatal de Investigación (AEI) and the Fondo Europeo de Desarrollo Regional (FEDER). JX was supported by the National Natural Science Foundation of China (Grant 32200682). NV was supported by funding from the German Research Foundation (DFG grants VE366/3-4 and VE366/12-1).

## Author contributions

AMV and JC conceived and designed the study.

MVV, PdlCR, DGG performed most *C. elegans* experiments.

EGG and JC performed RNAseq and qPCR.

MO performed luciferase assays.

MA and MJRP performed SeaHorse mitochondrial respiration assays.

JEI and NEB performed infection experiments.

NV, AS and SM performed lifespan experiments.

BM and JCC performed CelEST experiments.

JA, MM and RIF performed Ca^2+^ import measurements and electropharyngeograms.

WP, KB, MT and JP performed exopher analysis.

SX and JX performed laser wounding experiments.

AMV wrote the manuscript and all the authors edited and reviewed it.

## References

Alvarez-Illera, P., P. Garcia-Casas, J. Arias-Del-Val, R.I. Fonteriz, J. Alvarez et al., 2017 Pharynx mitochondrial [Ca(2+)] dynamics in live C. elegans worms during aging. Oncotarget 8 (34):55889–55900.

Alvarez-Illera, P., P. Garcia-Casas, R.I. Fonteriz, M. Montero, and J. Alvarez, 2020 Mitochondrial Ca(2+) Dynamics in MCU Knockout C. elegans Worms. Int J Mol Sci 21 (22).

Artal-Sanz, M., and N. Tavernarakis, 2009 Prohibitin couples diapause signalling to mitochondrial metabolism during ageing in C. elegans. Nature 461 (7265):793–797.

Artal-Sanz, M., W.Y. Tsang, E.M. Willems, L.A. Grivell, B.D. Lemire et al., 2003 The mitochondrial prohibitin complex is essential for embryonic viability and germline function in Caenorhabditis elegans. J Biol Chem 278 (34):32091–32099.

Banasiak, K., M. Turek, and W. Pokrzywa, 2023 Preparation of Caenorhabditis elegans for Scoring of Muscle-derived Exophers. Bio Protoc 13 (1).

Bast, A., and G.R. Haenen, 1988 Interplay between lipoic acid and glutathione in the protection against microsomal lipid peroxidation. Biochim Biophys Acta 963 (3):558–561.

Bennett, C.F., and M. Kaeberlein, 2014 The mitochondrial unfolded protein response and increased longevity: cause, consequence, or correlation? Exp Gerontol 56:142–146.

Berry, B.J., T.O. Nieves, and A.P. Wojtovich, 2021 Decreased Mitochondrial Membrane Potential Activates the Mitochondrial Unfolded Protein Response. MicroPubl Biol 2021.

Byrne, J.J., M.S. Soh, G. Chandhok, T. Vijayaraghavan, J.S. Teoh et al., 2019 Disruption of mitochondrial dynamics affects behaviour and lifespan in Caenorhabditis elegans. Cell Mol Life Sci 76 (10):1967–1985.

Cacho-Valadez, B., F. Munoz-Lobato, J.R. Pedrajas, J. Cabello, J.C. Fierro-Gonzalez et al., 2012 The characterization of the Caenorhabditis elegans mitochondrial thioredoxin system uncovers an unexpected protective role of thioredoxin reductase 2 in beta-amyloid peptide toxicity. Antioxid Redox Signal 16 (12):1384–1400.

Campbell, D., and S. Zuryn, 2024 The mechanisms and roles of mitochondrial dynamics in C. elegans. Semin Cell Dev Biol 156:266–275.

Charmpilas, N., and N. Tavernarakis, 2020 Mitochondrial maturation drives germline stem cell differentiation in Caenorhabditis elegans. Cell Death Differ 27 (2):601–617.

Cheng, Y.W., J. Liu, and T. Finkel, 2023 Mitohormesis. Cell Metab 35 (11):1872–1886.

Conrad, M., C. Jakupoglu, S.G. Moreno, S. Lippl, A. Banjac et al., 2004 Essential role for mitochondrial thioredoxin reductase in hematopoiesis, heart development, and heart function. Mol Cell Biol 24 (21):9414–9423.

Corsi, A.K., B. Wightman, and M. Chalfie, 2015 A Transparent Window into Biology: A Primer on Caenorhabditis elegans. Genetics 200 (2):387–407.

Curran, S.P., E.P. Leverich, C.M. Koehler, and P.L. Larsen, 2004 Defective mitochondrial protein translocation precludes normal Caenorhabditis elegans development. J Biol Chem 279 (52):54655–54662.

de Cubas, L., S. Boronat, M. Vega, A. Domenech, F. Gomez-Armengol et al., 2025 The glutathione system maintains the thiol redox balance in the mitochondria of fission yeast. Free Radic Biol Med 234:100–112.

de Lucas, M.P., A.G. Saez, and E. Lozano, 2015 miR-58 family and TGF-beta pathways regulate each other in Caenorhabditis elegans. Nucleic Acids Res 43 (20):9978–9993.

Dillin, A., A.L. Hsu, N. Arantes-Oliveira, J. Lehrer-Graiwer, H. Hsin et al., 2002 Rates of behavior and aging specified by mitochondrial function during development. Science 298 (5602):2398–2401.

Dong, Z., S. Shanmughapriya, D. Tomar, N. Siddiqui, S. Lynch et al., 2017 Mitochondrial Ca(2+) Uniporter Is a Mitochondrial Luminal Redox Sensor that Augments MCU Channel Activity. Mol Cell 65 (6):1014–1028 e1017.

Dorn, G.W., 2nd, 2019 Evolving Concepts of Mitochondrial Dynamics. Annu Rev Physiol 81:1–17.

Eriksson, S., J.R. Prigge, E.A. Talago, E.S. Arner, and E.E. Schmidt, 2015 Dietary methionine can sustain cytosolic redox homeostasis in the mouse liver. Nat Commun 6:6479.

Fiorese, C.J., A.M. Schulz, Y.F. Lin, N. Rosin, M.W. Pellegrino et al., 2016 The Transcription Factor ATF5 Mediates a Mammalian Mitochondrial UPR. Curr Biol 26 (15):2037–2043.

Flemming, A.J., Z.Z. Shen, A. Cunha, S.W. Emmons, and A.M. Leroi, 2000 Somatic polyploidization and cellular proliferation drive body size evolution in nematodes. Proc Natl Acad Sci U S A 97 (10):5285–5290.

Francis, R., and R.H. Waterston, 1991 Muscle cell attachment in Caenorhabditis elegans. J Cell Biol 114 (3):465–479.

Frokjaer-Jensen, C., M.W. Davis, C.E. Hopkins, B.J. Newman, J.M. Thummel et al., 2008 Single-copy insertion of transgenes in Caenorhabditis elegans. Nat Genet 40 (11):1375–1383.

Fu, H., H. Zhou, X. Yu, J. Xu, J. Zhou et al., 2020 Wounding triggers MIRO-1 dependent mitochondrial fragmentation that accelerates epidermal wound closure through oxidative signaling. Nat Commun 11 (1):1050.

Garbincius, J.F., and J.W. Elrod, 2022 Mitochondrial calcium exchange in physiology and disease. Physiol Rev 102 (2):893–992.

Gene Ontology, C., S.A. Aleksander, J. Balhoff, S. Carbon, J.M. Cherry et al., 2023 The Gene Ontology knowledgebase in 2023. Genetics 224 (1).

Gieseler, K., H. Qadota, and G.M. Benian, 2017 Development, structure, and maintenance of C. elegans body wall muscle. WormBook 2017:1–59.

Gomez-Orte, E., E. Cornes, A. Zheleva, B. Saenz-Narciso, M. de Toro et al., 2018 Effect of the diet type and temperature on the C. elegans transcriptome. Oncotarget 9 (11):9556–9571.

Gonzalez-Cabo, P., A. Bolinches-Amoros, J. Cabello, S. Ros, S. Moreno et al., 2011 Disruption of the ATP-binding cassette B7 (ABTM-1/ABCB7) induces oxidative stress and premature cell death in Caenorhabditis elegans. J Biol Chem 286 (24):21304–21314.

Gostimskaya, I., and C.M. Grant, 2016 Yeast mitochondrial glutathione is an essential antioxidant with mitochondrial thioredoxin providing a back-up system. Free Radic Biol Med 94:55–65.

Han, S.K., D. Lee, H. Lee, D. Kim, H.G. Son et al., 2016 OASIS 2: online application for survival analysis 2 with features for the analysis of maximal lifespan and healthspan in aging research. Oncotarget 7 (35):56147–56152.

Hanschmann, E.M., J.R. Godoy, C. Berndt, C. Hudemann, and C.H. Lillig, 2013 Thioredoxins, glutaredoxins, and peroxiredoxins--molecular mechanisms and health significance: from cofactors to antioxidants to redox signaling. Antioxid Redox Signal 19 (13):1539–1605.

Haynes, C.M., and S. Hekimi, 2022 Mitochondrial dysfunction, aging, and the mitochondrial unfolded protein response in Caenorhabditis elegans. Genetics 222 (4).

Jacobs, L., and J. Riemer, 2023 Maintenance of small molecule redox homeostasis in mitochondria. FEBS Lett 597 (2):205–223.

Jakupoglu, C., G.K. Przemeck, M. Schneider, S.G. Moreno, N. Mayr et al., 2005 Cytoplasmic thioredoxin reductase is essential for embryogenesis but dispensable for cardiac development. Mol Cell Biol 25 (5):1980–1988.

Kanehisa, M., M. Furumichi, Y. Sato, Y. Matsuura, and M. Ishiguro-Watanabe, 2025 KEGG: biological systems database as a model of the real world. Nucleic Acids Res 53 (D1):D672–D677.

Kelleher, J.F., M.A. Mandell, G. Moulder, K.L. Hill, S.W. L’Hernault et al., 2000 Myosin VI is required for asymmetric segregation of cellular components during C. elegans spermatogenesis. Curr Biol 10 (23):1489–1496.

Kim, E., A. Annibal, Y. Lee, H.H. Park, S. Ham et al., 2023 Mitochondrial aconitase suppresses immunity by modulating oxaloacetate and the mitochondrial unfolded protein response. Nat Commun 14 (1):3716.

Kim, H.E., A.R. Grant, M.S. Simic, R.A. Kohnz, D.K. Nomura et al., 2016 Lipid Biosynthesis Coordinates a Mitochondrial-to-Cytosolic Stress Response. Cell 166 (6):1539–1552 e1516.

Kramer, J.M., J.J. Johnson, R.S. Edgar, C. Basch, and S. Roberts, 1988 The sqt-1 gene of C. elegans encodes a collagen critical for organismal morphogenesis. Cell 55 (4):555–565.

Liu, Y., B.S. Samuel, P.C. Breen, and G. Ruvkun, 2014 Caenorhabditis elegans pathways that surveil and defend mitochondria. Nature 508 (7496):406–410.

Love, M.I., W. Huber, and S. Anders, 2014 Moderated estimation of fold change and dispersion for RNA-seq data with DESeq2. Genome Biol 15 (12):550.

Madaan, U., E. Yzeiraj, M. Meade, J.F. Clark, C.A. Rushlow et al., 2018 BMP Signaling Determines Body Size via Transcriptional Regulation of Collagen Genes in Caenorhabditis elegans. Genetics 210 (4):1355–1367.

Marty, L., D. Bausewein, C. Muller, S.A.K. Bangash, A. Moseler et al., 2019 Arabidopsis glutathione reductase 2 is indispensable in plastids, while mitochondrial glutathione is safeguarded by additional reduction and transport systems. New Phytol 224 (4):1569–1584.

Marty, L., W. Siala, M. Schwarzlander, M.D. Fricker, M. Wirtz et al., 2009 The NADPH-dependent thioredoxin system constitutes a functional backup for cytosolic glutathione reductase in Arabidopsis. Proc Natl Acad Sci U S A 106 (22):9109–9114.

Matsui, M., M. Oshima, H. Oshima, K. Takaku, T. Maruyama et al., 1996 Early embryonic lethality caused by targeted disruption of the mouse thioredoxin gene. Dev Biol 178 (1):179–185.

Mei, X., K.A. Maniates, A. Looper, A.R. Krauchunas, M. Druzhinina et al., 2023 SPE-51, a sperm-secreted protein with an immunoglobulin-like domain, is required for fertilization in C. elegans. Curr Biol 33 (14):3048–3055 e3046.

Melentijevic, I., M.L. Toth, M.L. Arnold, R.J. Guasp, G. Harinath et al., 2017 C. elegans neurons jettison protein aggregates and mitochondria under neurotoxic stress. Nature 542 (7641):367–371.

Melo, J.A., and G. Ruvkun, 2012 Inactivation of conserved C. elegans genes engages pathogen- and xenobiotic-associated defenses. Cell 149 (2):452–466.

Montero, M., M.T. Alonso, E. Carnicero, I. Cuchillo-Ibanez, A. Albillos et al., 2000 Chromaffin-cell stimulation triggers fast millimolar mitochondrial Ca2+ transients that modulate secretion. Nat Cell Biol 2 (2):57–61.

Monzel, A.S., J.A. Enriquez, and M. Picard, 2023 Multifaceted mitochondria: moving mitochondrial science beyond function and dysfunction. Nat Metab 5 (4):546–562.

Mora-Lorca, J.A., B. Saenz-Narciso, C.J. Gaffney, F.J. Naranjo-Galindo, J.R. Pedrajas et al., 2016 Glutathione reductase gsr-1 is an essential gene required for Caenorhabditis elegans early embryonic development. Free Radic Biol Med 96:446–461.

Muller, E.G., 1996 A glutathione reductase mutant of yeast accumulates high levels of oxidized glutathione and requires thioredoxin for growth. Mol Biol Cell 7 (11):1805–1813.

Munkacsy, E., M.H. Khan, R.K. Lane, M.B. Borror, J.H. Park et al., 2016 DLK-1, SEK-3 and PMK-3 Are Required for the Life Extension Induced by Mitochondrial Bioenergetic Disruption in C. elegans. PLoS Genet 12 (7):e1006133.

Naresh, N.U., and C.M. Haynes, 2019 Signaling and Regulation of the Mitochondrial Unfolded Protein Response. Cold Spring Harb Perspect Biol 11 (6).

Nargund, A.M., M.W. Pellegrino, C.J. Fiorese, B.M. Baker, and C.M. Haynes, 2012 Mitochondrial import efficiency of ATFS-1 regulates mitochondrial UPR activation. Science 337 (6094):587–590.

Nicolas-Avila, J.A., A.V. Lechuga-Vieco, L. Esteban-Martinez, M. Sanchez-Diaz, E. Diaz-Garcia et al., 2020 A Network of Macrophages Supports Mitochondrial Homeostasis in the Heart. Cell 183 (1):94–109 e123.

Nonn, L., R.R. Williams, R.P. Erickson, and G. Powis, 2003 The absence of mitochondrial thioredoxin 2 causes massive apoptosis, exencephaly, and early embryonic lethality in homozygous mice. Mol Cell Biol 23 (3):916–922.

Nystrom, J., Z.Z. Shen, M. Aili, A.J. Flemming, A. Leroi et al., 2002 Increased or decreased levels of Caenorhabditis elegans lon-3, a gene encoding a collagen, cause reciprocal changes in body length. Genetics 161 (1):83–97.

Olmedo, M., M. Geibel, M. Artal-Sanz, and M. Merrow, 2015 A High-Throughput Method for the Analysis of Larval Developmental Phenotypes in Caenorhabditis elegans. Genetics 201 (2):443–448.

Page, A.P., and I.L. Johnstone, 2007 The cuticle. WormBook:1–15.

Palikaras, K., E. Lionaki, and N. Tavernarakis, 2015 Coordination of mitophagy and mitochondrial biogenesis during ageing in C. elegans. Nature 521 (7553):525–528.

Palikaras, K., and N. Tavernarakis, 2016 Intracellular Assessment of ATP Levels in Caenorhabditis elegans. Bio Protoc 6 (23).

Picca, A., J. Faitg, J. Auwerx, L. Ferrucci, and D. D’Amico, 2023 Mitophagy in human health, ageing and disease. Nat Metab 5 (12):2047–2061.

Porras, P., J.R. Pedrajas, E. Martinez-Galisteo, C.A. Padilla, C. Johansson et al., 2002 Glutaredoxins catalyze the reduction of glutathione by dihydrolipoamide with high efficiency. Biochem Biophys Res Commun 295 (5):1046–1051.

Powell, J.R., and F.M. Ausubel, 2008 Models of Caenorhabditis elegans infection by bacterial and fungal pathogens. Methods Mol Biol 415:403–427.

Prigge, J.R., S. Eriksson, S.V. Iverson, T.A. Meade, M.R. Capecchi et al., 2012 Hepatocyte DNA replication in growing liver requires either glutathione or a single allele of txnrd1. Free Radic Biol Med 52 (4):803–810.

Pujol, N., S. Cypowyj, K. Ziegler, A. Millet, A. Astrain et al., 2008 Distinct innate immune responses to infection and wounding in the C. elegans epidermis. Curr Biol 18 (7):481–489.

Raizen, D.M., R.Y. Lee, and L. Avery, 1995 Interacting genes required for pharyngeal excitation by motor neuron MC in Caenorhabditis elegans. Genetics 141 (4):1365–1382.

Rea, S.L., N. Ventura, and T.E. Johnson, 2007 Relationship between mitochondrial electron transport chain dysfunction, development, and life extension in Caenorhabditis elegans. PLoS Biol 5 (10):e259.

Restif, C., C. Ibanez-Ventoso, M.M. Vora, S. Guo, D. Metaxas et al., 2014 CeleST: computer vision software for quantitative analysis of C. elegans swim behavior reveals novel features of locomotion. PLoS Comput Biol 10 (7):e1003702.

Robinson, M.D., D.J. McCarthy, and G.K. Smyth, 2010 edgeR: a Bioconductor package for differential expression analysis of digital gene expression data. Bioinformatics 26 (1):139–140.

Rodriguez-Palero, M.J., A. Lopez-Diaz, R. Marsac, J.E. Gomes, M. Olmedo et al., 2018 An automated method for the analysis of food intake behaviour in Caenorhabditis elegans. Sci Rep 8 (1):3633.

Rogers, L.K., T. Tamura, B.J. Rogers, S.E. Welty, T.N. Hansen et al., 2004 Analyses of glutathione reductase hypomorphic mice indicate a genetic knockout. Toxicol Sci 82 (2):367–373.

Runkel, E.D., R. Baumeister, and E. Schulze, 2014 Mitochondrial stress: balancing friend and foe. Exp Gerontol 56:194–201.

Schiavi, A., S. Maglioni, K. Palikaras, A. Shaik, F. Strappazzon et al., 2015 Iron-Starvation-Induced Mitophagy Mediates Lifespan Extension upon Mitochondrial Stress in C. elegans. Curr Biol 25 (14):1810–1822.

Schiavi, A., E. Salveridou, V. Brinkmann, A. Shaik, R. Menzel et al., 2023 Mitochondria hormesis delays aging and associated diseases in Caenorhabditis elegans impacting on key ferroptosis players. iScience 26 (4):106448.

Schneider, C.A., W.S. Rasband, and K.W. Eliceiri, 2012 NIH Image to ImageJ: 25 years of image analysis. Nat Methods 9 (7):671–675.

Shpilka, T., Y. Du, Q. Yang, A. Melber, N. Uma Naresh et al., 2021 UPR(mt) scales mitochondrial network expansion with protein synthesis via mitochondrial import in Caenorhabditis elegans. Nat Commun 12 (1):479.

Shpilka, T., and C.M. Haynes, 2018 The mitochondrial UPR: mechanisms, physiological functions and implications in ageing. Nat Rev Mol Cell Biol 19 (2):109–120.

Sies, H., C. Berndt, and D.P. Jones, 2017 Oxidative Stress. Annu Rev Biochem 86:715–748.

Sies, H., R.J. Mailloux, and U. Jakob, 2024 Fundamentals of redox regulation in biology. Nat Rev Mol Cell Biol 25 (9):701–719.

Soo, S.K., and J.M. Van Raamsdonk, 2021 High confidence ATFS-1 target genes for quantifying activation of the mitochondrial unfolded protein response. MicroPubl Biol 2021.

Stenvall, J., J.C. Fierro-Gonzalez, P. Swoboda, K. Saamarthy, Q. Cheng et al., 2011 Selenoprotein TRXR-1 and GSR-1 are essential for removal of old cuticle during molting in Caenorhabditis elegans. Proc Natl Acad Sci U S A 108 (3):1064–1069.

Stiernagle, T., 2006 Maintenance of C. elegans. WormBook:1–11.

Suzuki, Y., G.A. Morris, M. Han, and W.B. Wood, 2002 A cuticle collagen encoded by the lon-3 gene may be a target of TGF-beta signaling in determining Caenorhabditis elegans body shape. Genetics 162 (4):1631–1639.

Tabara, L.C., M. Segawa, and J. Prudent, 2025 Molecular mechanisms of mitochondrial dynamics. Nat Rev Mol Cell Biol 26 (2):123–146.

Tan, S.X., D. Greetham, S. Raeth, C.M. Grant, I.W. Dawes et al., 2010 The thioredoxin-thioredoxin reductase system can function in vivo as an alternative system to reduce oxidized glutathione in Saccharomyces cerevisiae. J Biol Chem 285 (9):6118–6126.

Teoh, J.S., M.S. Soh, J.J. Byrne, and B. Neumann, 2019 Quantitative Approaches for Studying Cellular Structures and Organelle Morphology in Caenorhabditis elegans. J Vis Exp (149).

Tjahjono, E., A.P. McAnena, and N.V. Kirienko, 2020 The evolutionarily conserved ESRE stress response network is activated by ROS and mitochondrial damage. BMC Biol 18 (1):74.

Traa, A., A. Keil, A. AlOkda, S. Jacob-Tomas, A.A. Tamez Gonzalez et al., 2024 Overexpression of mitochondrial fission or mitochondrial fusion genes enhances resilience and extends longevity. Aging Cell 23 (10):e14262.

Turek, M., K. Banasiak, M. Piechota, N. Shanmugam, M. Macias et al., 2021 Muscle-derived exophers promote reproductive fitness. EMBO Rep 22 (8):e52071.

Ventura, N., S. Rea, S.T. Henderson, I. Condo, T.E. Johnson et al., 2005 Reduced expression of frataxin extends the lifespan of Caenorhabditis elegans. Aging Cell 4 (2):109–112.

Wang, E., Y. Jiang, and C. Zhao, 2024a Structural and physiological functions of Caenorhabditis elegans epidermis. Heliyon 10 (19):e38680.

Wang, L., W. Li, X. Wu, Q. Ouyang, B. Sun et al., 2025 FGF21 maintains redox homeostasis and promotes neuronal survival after traumatic brain injury by targeting SLC25A39-mediated mitochondrial GSH transport. J Transl Med 23 (1):1044.

Wang, P., L. Chen, N. Wang, L. Miao, and Y. Zhao, 2024b Mitochondrial defects triggered by amg-1 mutation elicit UPRmt and phagocytic clearance during spermatogenesis in C. elegans. Development 151 (3).

Wang, Y., F.S. Yen, X.G. Zhu, R.C. Timson, R. Weber et al., 2021 SLC25A39 is necessary for mitochondrial glutathione import in mammalian cells. Nature 599 (7883):136–140.

Willems, P.H., R. Rossignol, C.E. Dieteren, M.P. Murphy, and W.J. Koopman, 2015 Redox Homeostasis and Mitochondrial Dynamics. Cell Metab 22 (2):207–218.

Xu, S., and A.D. Chisholm, 2014a C. elegans epidermal wounding induces a mitochondrial ROS burst that promotes wound repair. Dev Cell 31 (1):48–60.

Xu, S., and A.D. Chisholm, 2014b Methods for skin wounding and assays for wound responses in C. elegans. J Vis Exp (94).

Yang, W., and S. Hekimi, 2010 Two modes of mitochondrial dysfunction lead independently to lifespan extension in Caenorhabditis elegans. Aging Cell 9 (3):433–447.

Yoneda, T., C. Benedetti, F. Urano, S.G. Clark, H.P. Harding et al., 2004 Compartment-specific perturbation of protein handling activates genes encoding mitochondrial chaperones. J Cell Sci 117 (Pt 18):4055–4066.

Zhang, H., X. Li, W. Fan, S. Pandovski, Y. Tian et al., 2023 Inter-tissue communication of mitochondrial stress and metabolic health. Life Metab 2 (1).

Zhao, Y.G., Y. Chen, G. Miao, H. Zhao, W. Qu et al., 2017 The ER-Localized Transmembrane Protein EPG-3/VMP1 Regulates SERCA Activity to Control ER-Isolation Membrane Contacts for Autophagosome Formation. Mol Cell 67 (6):974–989 e976.

Zhou, L., and Y. Liu, 2025 The trade-off between reproduction and resilience. Trends Endocrinol Metab.

